# Satellite-gene abundance fusion reveals global hotspots of ocean ecosystem services

**DOI:** 10.64898/2026.09.23.753815

**Authors:** Baptiste Serandour, Constanza M. Andreani-Gerard, Antoine Bagnaro, Sacha Bourgeois-Gironde, Samuel Chaffron, Andrés Couve, Roy El Hourany, Lionel Guidi, Iñaki Hojas, Andre Abreu, Alejandro Maass, Damien Eveillard

## Abstract

The Ocean sustains planetary health through climate regulation, biodiversity, and essential ecosystem services. Plankton underpins these processes by driving primary production, sustaining food webs, and mediating carbon sequestration. Yet the spatial organization of plankton-based ecosystem services remains poorly understood. The High Seas Treaty and the 30% Ocean protection target by 2030 create an urgent need for science-based guidance for global ocean governance. Here, we aim to identify key planktonic areas associated with functional diversity, net primary productivity, and carbon export globally. We used machine learning to integrate omics-derived plankton functions and satellite observations with diversity metric and ecosystem services. Leveraging metagenomic data, we identified molecular functions strongly linked to ecosystem functioning and project their global distributions across seasons. We reveal putative hotspots of functional diversity, net primary productivity, and carbon export. Our analysis pinpoints areas where all three indicators converge synchronously, predominantly along oceanographic structures such as polar fronts and principal current systems. By relying on satellite data as core input, our approach captures the temporal variability of these hotspots, providing a global, dynamic protocol for projections of plankton-based ecosystem services. These findings establish a foundation for developing integrative conservation tools based on ecosystem functioning and inform adaptive management under climate change.

## Introduction

Despite its importance, the ocean is increasingly threatened by climate change and other anthropogenic pressures, raising concerns about the long-term functioning and resilience of marine ecosystems [1, 2]. In response, the international community has committed to ambitious conservation targets. The “30 by 30” initiative was adopted in 2022 by more than 190 countries. It is a global conservation project aimed at protecting 30% of the Earth’s oceans, freshwater, and land by 2030. In addition, the High Seas Treaty (also known as the Biodiversity Beyond National Jurisdiction, BBNJ) is a United Nations program and a legally binding international treaty that seeks to protect international waters. To achieve these goals and effectively protect the ocean, global science-based tools are necessary to identify key marine regions that deliver the greatest ecological and climatic benefits.

Among conservation tools, Marine Protected Areas (MPAs) are the most widely implemented, and their number and surface area have expanded rapidly [3]. MPAs are designed to protect specific species or habitats by reducing degradation from human activities [4]. These ocean conservation measures have largely been focused on indicators of fish biodiversity or resources, such as species richness or habitat extent, and have mostly been applied to national waters [3]. Indeed, as of 2026, only 1.45% of areas beyond national jurisdiction (ABNJ) have been protected, even though they represent 61% of the global surface ocean [5]. Part of this difference is attributable to a lack of knowledge about high-seas ecosystems and their management needs [3]. Over the past decades, ABNJ have witnessed an exponential growth in human activities, including transport and fishing, which raise concerns about the overexploitation of fish stocks [6]. Although MPAs are recognized as aiming to sustain ocean services and being efficient for marine species protection [7], their distribution is not optimal for ecosystem services [8]. In particular, the static approach of MPAs does not align with the dynamic nature of the pelagic component of the ocean [8], as its circulation generates transient features such as fronts, filaments, and eddies that persist from weeks to months, creating pronounced spatial and temporal heterogeneity in biogeochemical processes [9, 10]. Moreover, climate change is altering the intensity, frequency, and spatial distribution of these processes, leading to ongoing shifts in species distributions and ecosystem functioning that are projected to continue throughout the coming decades [10–12].

The planktonic part of the ocean plays an essential but overlooked role in delivering ecosystem services that control climate [13, 14], support major biogeochemical cycles [15, 16], sustain marine food [17] and the global economy [18]. Through the biological carbon pump (BCP), marine plankton communities export and trap organic carbon, thereby removing an estimated 2.81 Gt of carbon from the atmosphere each year [19], thereby significantly mitigating the impacts of climate change. In parallel, marine planktonic ecosystems host extraordinary and largely unexplored biodiversity, with estimates suggesting that up to 90% of marine species remain undiscovered [20]. Together, these functions and the biodiversity they support make marine plankton a fundamental component of ocean functioning, underpinning the ecological, climatic, and economic services provided by the ocean. Thus, the dynamic plankton-mediated processes that underpin key ecosystem services, including carbon export and primary production, remain largely absent from marine spatial planning. For instance, recent work suggests that most of the carbon sequestration by the BCP within economic exclusive zones occurs outside MPAs [19], highlighting the need for improved tools to identify ocean regions that sustain critical ecosystem functions. This emphasizes the need for new protocols that can integrate cutting-edge omics data to explicitly bring plankton-derived processes into marine conservation and spatial planning frameworks.

Tracking plankton dynamics is essential to better understand marine community shifts. Satellite remote sensing is an efficient way to do so, as it provides global coverage within short time-frames and enables the measurement of key parameters of plankton distribution [21]. Most studies linking satellites to marine biological components have focused on phytoplankton, as pigment production serves as a proxy for the phototrophic community [22, 23]. Recent work has taken steps toward a more complete understanding of plankton communities notably by predicting size classes [24] or even by combining omics data from phytoplankton and heterotrophic protists with Earth observations to predict community types [25]. However, these techniques do not benefit from advances in marine metagenomics, which have opened an unprecedented window into the functional potential of plankton communities across the global ocean [26]. Notably, metagenomic data capture the metabolic capabilities of resident microbial communities, offering a functional perspective relevant for the BCP. Despite this transformative potential, the richness of functional information from global plankton genomic surveys has not yet been fully integrated into marine conservation planning. Resolving how this functional potential varies across space and time would provide a more complete picture of plankton-driven ecosystem functioning and services. Beyond advancing our understanding of pelagic ecosystem dynamics, such knowledge could help address the long-recognized mismatch between static conservation measures and dynamic pelagic ecosystems [27, 28].

To do so, we developed a *Key Ocean Planktonic Areas* (KOPAs) protocol, a science-based and data-driven modelling approach designed to identify ocean regions that provide high levels of plankton-mediated ecosystem services. Our KOPAs protocol leverages the seminal high-resolution metagenomic plankton survey collected during major oceanographic expeditions (i.e., Tara Oceans, Tara Polar Circle, Malaspina, Bio-GO-SHIP and GEOTRACES, see Figure S1), which collectively represent one of the most comprehensive efforts to genetically characterize marine microbial life. These molecular datasets were processed to identify specific genetic prokaryotic functions associated with essential ecosystem states, including carbon export, net primary production (NPP), and a functional diversity metric computed as the Shannon index of genome-resolved abundance profiles (hereafter functional diversity). The KOPAs protocol further combines them with remote sensing observations, using machine learning approaches, to project the surface distribution of these functions with the goal to forecast the spatial distribution of a new generation of omics-informed ecosystem metrics. Carbon export, NPP, and functional diversity are key ecological indicators of ocean ecosystem functioning, as they together reflect the efficiency of oceanic biological carbon pump, the energetic capacity of the base of marine food webs [29, 30], as well as the structural complexity and resilience of planktonic communities. Thus, the KOPAs protocol enables several levels of application for the understanding of the marine environment. First, it can be used to assess the current MPAs design’s ability to capture plankton-based ecosystem services and functional diversity. Second, it may be useful to monitor the distribution of molecular functions across oceans and to provide virtual projection-based sampling alongside *in-situ* observations, as this represents a frugal, time-efficient complement to oceanographic cruise sampling. Finally, the KOPAs protocol identifies priority regions for conservation programs based on plankton-derived ecosystem services by dynamically tracking emerging areas of ecological importance driven by plankton functional potential.

### KOPAs protocol

The KOPAs protocol comprises three sequential steps (see Figure 1). First, we identify KEGG Orthologs (KOs), which group genes with conserved functions and provide a standardized representation of molecular functions and metabolic pathways, derived from procaryotic MAGs associated with each observed ecological indicator (NPP, carbon export and functional diversity) through a two-stage filtering procedure applied across all samples. Initially, a weighted gene co-expression network analysis (WGCNA, [31]) was performed using the *ρ* proportionality metric to minimize spurious correlations arising from the compositional nature of omics data [32]. The WGCNA creates modules of KOs with similar abundance patterns. The modules significantly associated with each observed ecological indicator were identified, yielding three indicator-specific KO subsets. The subsets associated with NPP and functional diversity were subsequently refined using Variable Importance in Projection (VIP) scores from partial least squares regression (PLSr). In the second step, we assessed the predictability of individual normalized abundances of KOs to identify those that could be estimated (*R*^2^ *>* 0.3 for NPP and functional diversity; *R*^2^ *>* 0.2 for carbon export) from remote sensing observations. Namely, environmental variables obtained from satellite data comprised backscattering at 443nm (bbp443), photosynthetically available radiation (PAR), total photosynthetically available radiation (IPAR), sea surface temperature (SST), Chlorophyll a, and the diffuse attenuation coefficient for downwelling irradiance at 490 nm (kd490). These selected satellite variables are broadly consistent with the main components represented in bio-optical NPP models: phytoplankton biomass or pigmentation through chlorophyll-a and bbp443, light availability and its vertical attenuation through PAR, iPAR, and kd490, and environmental regulation of phytoplankton physiology through SST. XGBoost [33] was selected for its consistently highest predictive performance. As a third step, we built and trained a second machine learning model using observed KO abundances as input to quantify the predictive power of the selected KOs for the target ecological indicator (Figure 2A). We then applied the full pipeline to global-scale satellite observations to generate global predictions of KO abundance by using the first model, which were subsequently fed into the second model to project the spatial distribution of each ecological indicator across the year 2023 (Figure 2B–D). Ecological indicators hotspots were then compared with the distribution of marine protected areas [34].

**Fig. 1.**
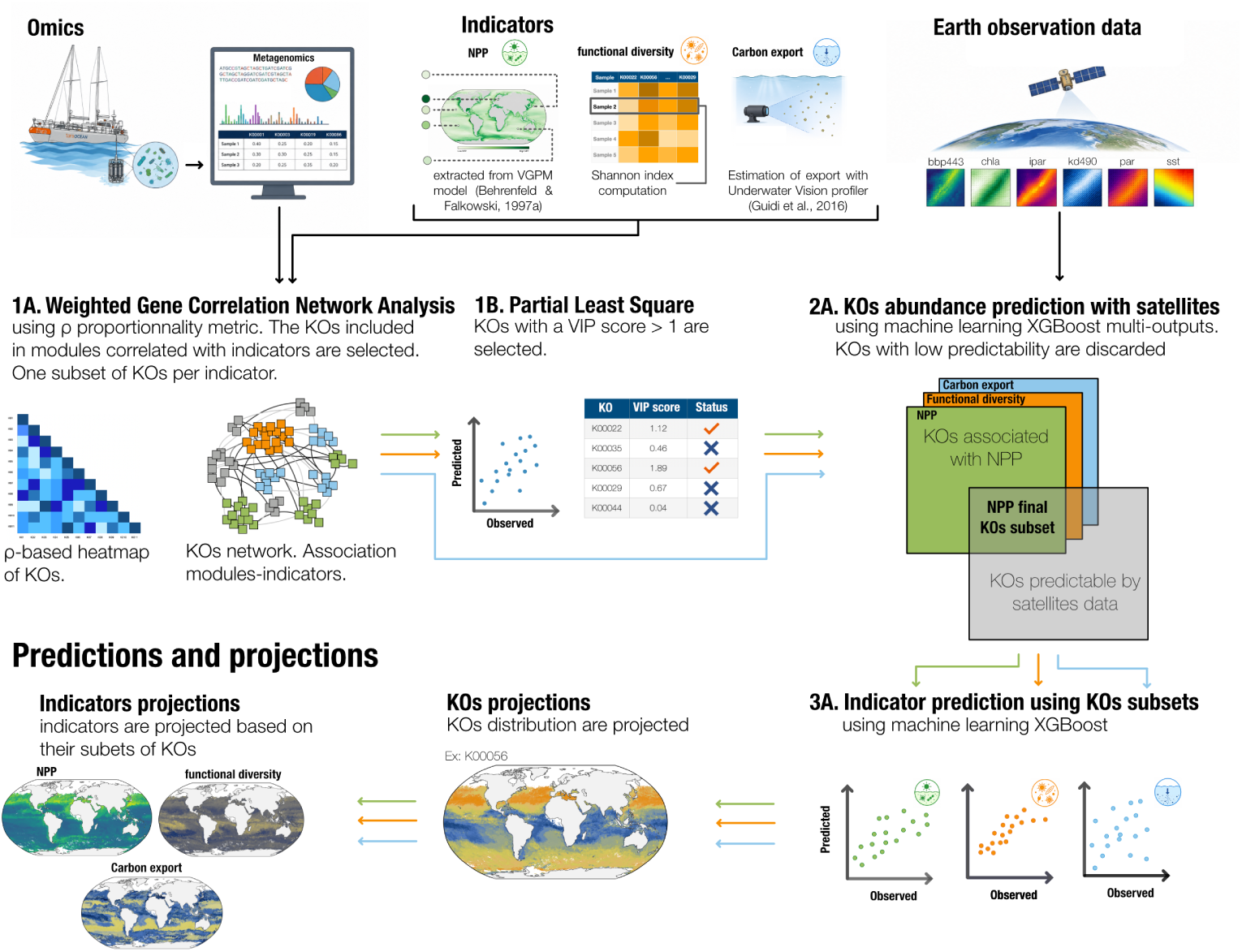
Rationale of the KOPAs protocol that transforms omics knowledge into ecological indicators distribution. The KOPAs protocol comprises three steps. First, for each indicator (carbon export, NPP, functional diversity), we performed network analysis (i.e., *ρ*-based WGCNA) to identify correlated KO modules(*|r| >* 0.3, *p <* 0.05; 1A). A module embeds a set of KOs that altogether is a good predictor for a given indicator. Second, we applied PLS regression on the NPP and functional diversity subsets, retaining features with VIP scores above the threshold (1B). Third, we filtered KOs based on their predictability using satellite data through an Extreme Gradient Boosting (XGBoost) model (2A). The final KO subset for each indicator was then used to predict indicator values (3A), followed by global projections mapping of predicted KOs abundances from which predicted indicator values are estimated (see Methods for details).

**Fig. 2.**
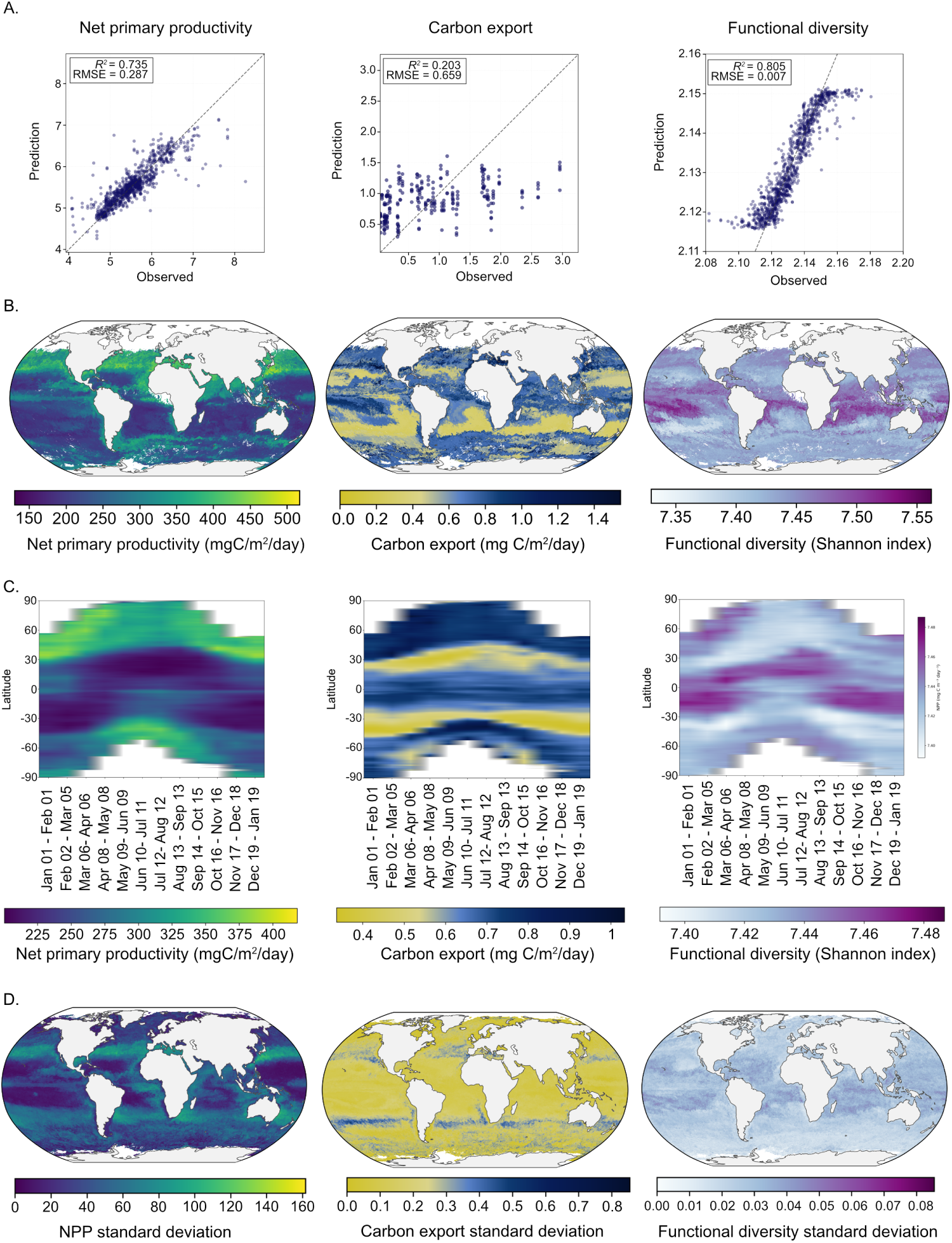
Indicator prediction and spatio-temporal variability of the projection. A. Predicted values are derived from a selected subset of KOs: the ability of KOs abundance profiles subsets to predict indicator values. B. Global projection of the ecological indicators for the period ranging from Jan 01 to 01 Feb 2023 using the KOPAs framework. C. Temporal variability of the predicted indicators across longitudes and months during the year 2023. D. Standard deviation of the predicted indicators on global projections across months during the year 2023.

### KOs selection and metabolic validation

We performed a two-step filtration process on KOs to select those most closely related to our indicators (Figure 1). The WGCNA step induces the selection of six modules for NPP (1360 KOs), ten for functional diversity (4263 KOs), and one for carbon export (106 KOs). Following the selection of KOs using WGCNA, we used PLS-r for VIP-based selection (see Method section), yielding subsets of 531 and 1773 KOs for NPP and functional diversity, respectively. No additional selection of KOs associated to carbon export was performed due to the PLSr’s low reliability.

To investigate the predictability of KOs using environmental variables from satellites, we developed an XGBoost multi-output model (see Figure S2 for variable importance). Among the NPP subset, 462 KOs were predictable using satellites with an *R*^2^ *>*0.3, and 1197 from the functional diversity subset met the same criterion. Due to the limited amount of data, an *R*^2^ *>*0.2 filter was applied to carbon export, resulting in the selection of 42 KOs. The remaining KOs after these filtering steps were used for projections (Figure S3).

Annotations of metabolic pathways for selected KOs were analyzed for biological validation. Our results show that 176 (38%), 481 (40%), and 20 (48%) KOs associated with NPP, functional diversity, and carbon export, respectively, were of unknown function. To ensure biological interpretability, the functional characterization of each ecological indicator was based solely on the subsets of KOs that had annotated pathways (Figure 3). We observe that 30% of NPP pathways are involved in energy metabolism. Remarkably, two-thirds of those pathways are linked to photosynthesis and about one-fifth to oxidative phosphorylation. Porphyrin and terpenoid-quinone metabolism stood out among other cofactors and vitamins (Figure 3 and S4). While porphyrin rings constitute chlorophylls, light-harvesting structures, ubiquinones and plastoquinones are electron carriers that shuttle within photosynthetic and respiratory membranes. Photosynthesis and respiration, two fundamental bioenergetic processes act as twin engines of the pelagic ecosystem: the first fixes solar energy into reduced organic carbon; the second liberates this stored chemical energy to power cellular maintenance and growth [35, 36].

**Fig. 3.**
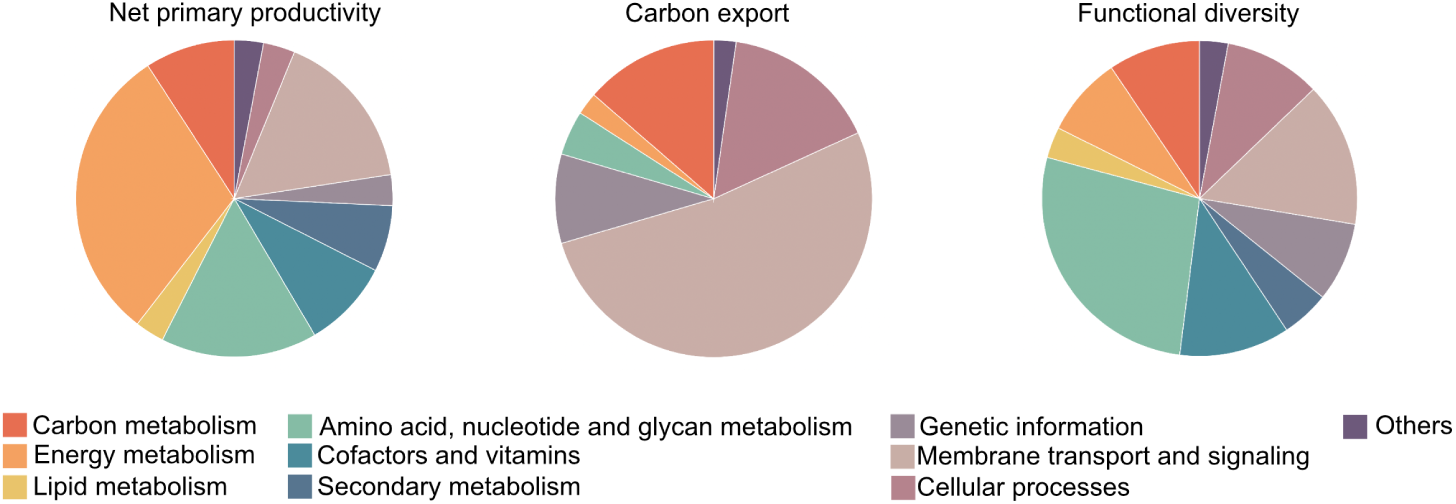
Metabolic characterization of selected KOs. Distribution of metabolic pathways annotated for each subset of KOs associated with an ecological indicator. Functional categories follow the KEGG nomenclature.

For functional diversity, 27% of pathways are involved in amino acid, nucleotide, and glycan metabolism, with a major contribution of aromatic amino acids, i.e., phenylalanine, tyrosine, and tryptophan. Biofilm formation stood out compared to other cellular processes (Figures 3 and S4). Nucleotides encode for genes and amino acids are the translated function, compiling a community’s functional capacity to build proteins. Aromatic amino acids are precursors to many bioactive compounds, including signaling molecules and secondary metabolites. They capture refined mechanisms of microbial interaction and adaptation to changing environments [37, 38].

Lastly, our results indicate that more than half of the pathways retrieved for carbon export are involved in membrane transport and signaling; namely, bacterial secretion systems (27%), two-component systems (16%), and ABC transporters (9%), all of which participate or enable the active release of not only extracellular polymeric substances, but also hydrolytic enzymes. Some have the potential to aggregate and affect the sinking speed of particulate organic carbon, while others degrade organic matter and increase the lability of dissolved organic carbon [39, 40]. Cellular processes comprising cell growth, death, and quorum sensing accounted for 11% of pathways, while replication and repair accounted for 9% (Figures 3 and S4). At the microbial community level, these functions may regulate carbon secretion in response to population density and environmental cues [41, 42].

### Ecological indicators reconstruction

Using the observed abundance of the above KO selection and satellite predictability, we developed a machine learning model to estimate the ability of our subset of KOs to predict the log(1p)-transformed NPP, carbon export and functional diversity levels (Figure 2). We observed a correlation of 0.735 between predicted and NPP values from Behrenfeld Falkowski, 1997 [43], with an RMSE of 0.287. For functional diversity, we obtained an *R*^2^ of 0.805 and an RMSE of 0.007, whereas for carbon export, the *R*^2^ was 0.203 and the RMSE was 0.659 (Figure 2A). Additionally, we performed a Spearman rank correlation that revealed a significant positive relationship between the observed and predicted carbon export values (Spearman’s rank correlation coefficient = 0.496, n = 51, p-value *<* 0.01), indicating that the KOPAs framework captures a meaningful fraction of the qualitative variability in carbon export. The predictive performance of our KOPAS protocol is consistent with existing satellite-based approaches. In particular, the performance of our global NPP model is comparable to recent omics-based predictions (*R*^2^ = 0.83 for chlorophyll-a vs. *in-situ* observations [26]). Critically, by routing predictions through a molecular pathway intermediate, the KOPAs protocol captures the likely molecular processes that drive productivity rather than their optical signatures.

### Methodological validation

To further evaluate the robustness of the KOPAs protocol at a global scale, we assessed the reliability of its predictions of ecological indicator. As a consistency assessment with the reference NPP product used for model development, estimates of NPP can be compared with established satellite-derived NPP products, thereby methodologically validating the protocol beyond the standard training dataset. Although the KOPAs protocol reproduces the global spatial patterns of the VGPM model, its projections exhibit substantially lower magnitudes in high-NPP environments, such as coastal shelves, upwelling areas, and frontal systems (Figure 4). Projections of NPP-based key areas using Carbon-based Productivity Model (CbPM, [44]) and Carbon, Absorption, and Fluorescence Euphotic-resolving (CAFE, [45]), two alternative sources from NPP estimates were compared to the KOPAs outputs(Supplementary material 10). The uses of these models exhibited substantial spatial overlap with VGPM-derived patterns (ranging from 57.6–71.9% for CBPM and 21.4–34.8% for CAFE – Figure S5).

**Fig. 4.**
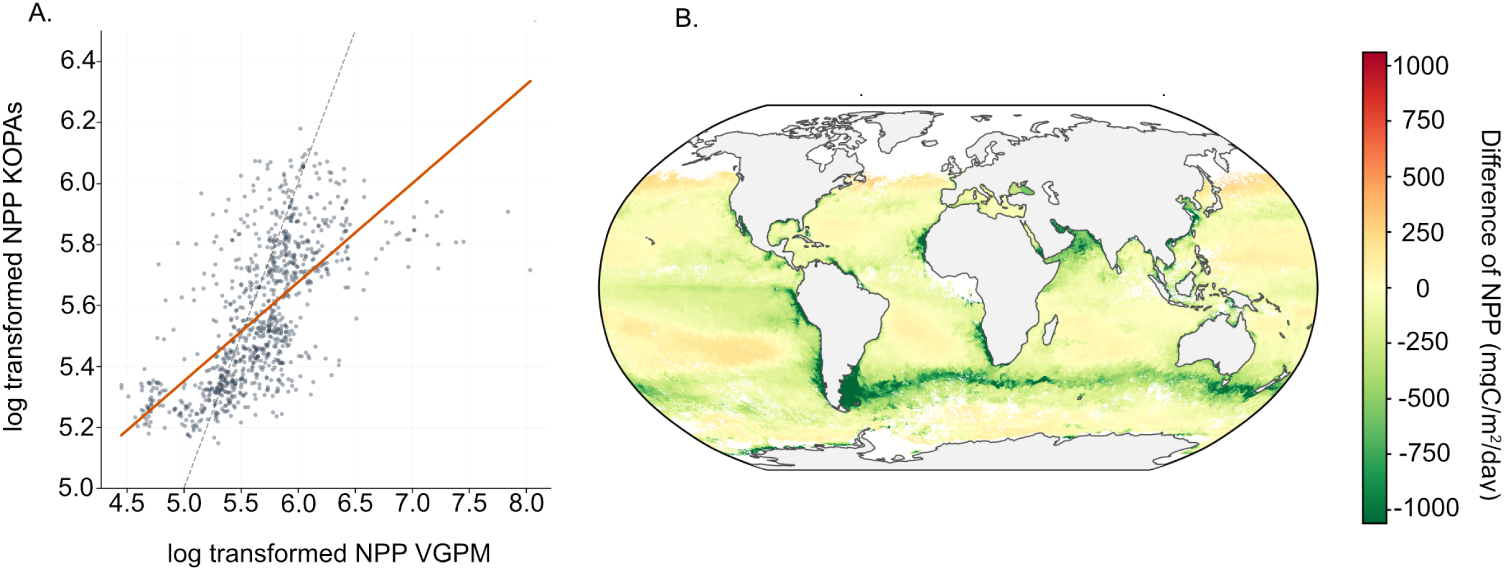
Comparison of the NPP outputs from the KOPAs protocol. A. Comparison of the KOPAs value with the seminal Vertically Generalized Production Model (VGPM; [43]) values at the *in-situ* omics sampling locations. B. Difference between KOPAs protocol-based NPP prediction as shown in 2B, and VGPM as implemented by the Ocean Productivity site for the same period. Zero values (i.e., yellow) indicate no differences between the two models. Positive values (i.e., red color code) indicate higher NPP estimates in the KOPAs protocol than in VGPM, and negative values (i.e., green color code) indicate the opposite.

These KOPAs outputs are reinforced by the oceanographic coherence of our projections. We delineate areas of triple concurrence of NPP, carbon export and functional diversity at high intensities—designated here as KOPAs (Figure 5). Triple concurrence refers to locations where NPP, carbon export, and functional diversity simultaneously exceed their respective high-value thresholds during the same temporal period. These KOPAs display a latitudinal symmetry about 40° and undergo substantial seasonal reorganization across the monthly projection during the year 2023 (Figure S9), reflecting the dynamic nature of plankton-based ecosystem services. Rather than occurring uniformly, they map preferentially onto major oceanographic discontinuities such as the Gulf Stream’s mesoscale fronts (from September to October), the Subtropical Front of the Antarctic Circumpolar Current, as well as the northern Atlantic and Pacific basins during the boreal winter. Regionally, the Costa Rica thermal dome (from February to April), upwelling off Chile (from April to August) and off Senegal, together with wintertime signals across the Mediterranean, were identified as KOPAs. The spatial distribution of high-value plankton-based ecosystem services varies substantially across months (Figure 2B and 2D). The ecological processes underpinning these indicators are therefore characterized not only by spatial heterogeneity but also by temporal variability (Figure 2B and S9). These findings are particularly relevant for pelagic systems, where ecological significance is often associated with biological activity rather than with stable physical features. The results provide empirical support for management frameworks that incorporate periodic reassessment of protected areas and spatially dynamic conservation measures, informed by updated ecological information.

**Fig. 5.**
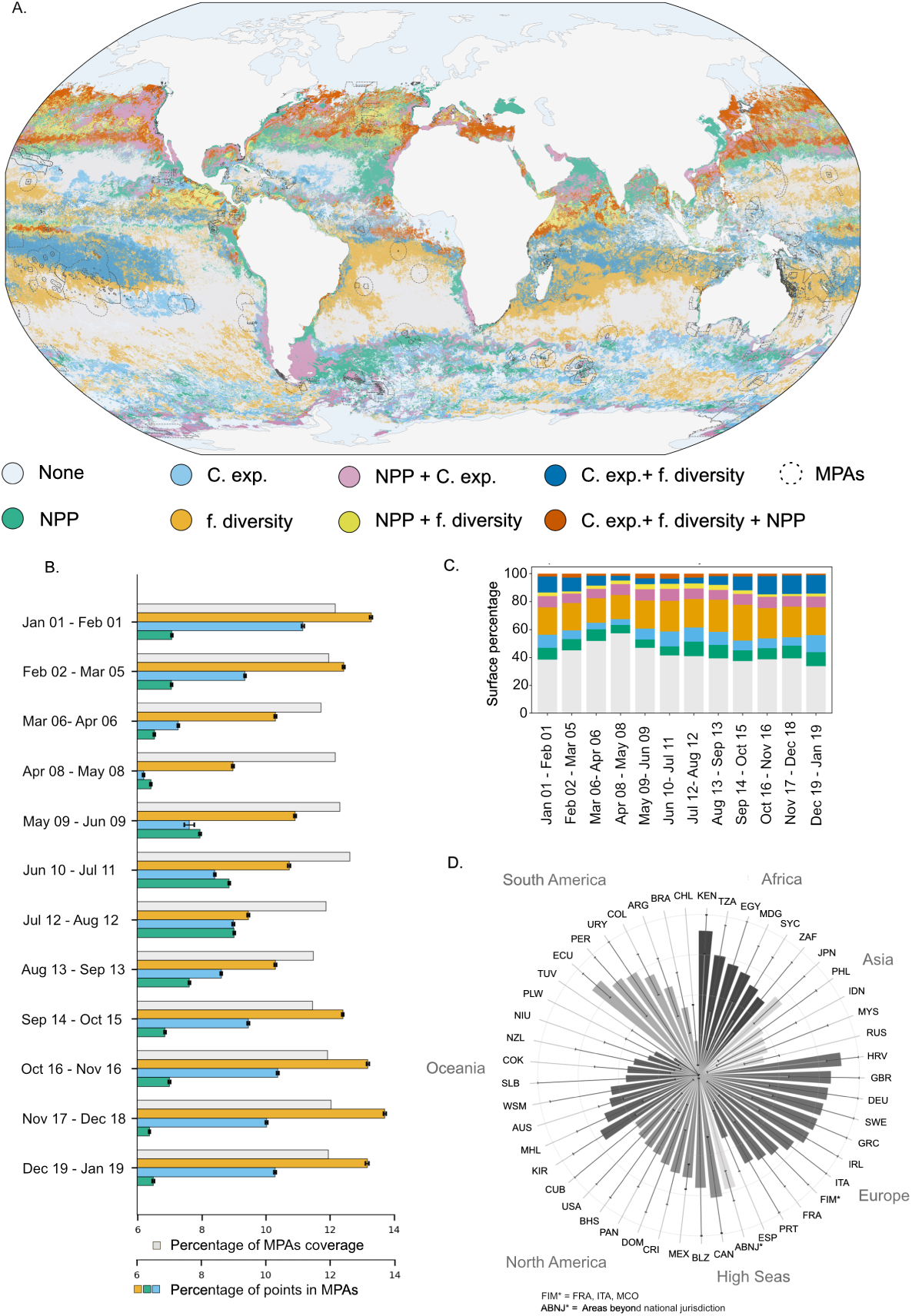
Identification of key ocean planktonic areas. A. Global distribution of projected indicator combinations for the period 12 July – 12 August 2023. B. Comparison of the proportion of ocean covered by MPAs with the proportion of high indicator levels within MPAs, the error bars show the variability across 60 subsamplings. C. Temporal variability in the monthly coverage of MPAs by individual indicators or their combinations. D. Analysis per country of the proportion of MPAs surface covered by at least one high-value indicator, the black error bars show the q10 and q90 of the temporal variability across the year 2023.

A key strength of the KOPAs protocol lies in its explainability. Projections of ecological indicators emerge from the collective distribution and equilibrium of projected omics-derived features, enabling variability in indicator levels to be traced directly to underlying molecular variation. By focusing on the distribution of molecular functions rather than taxonomic units, the KOPAs protocol circumvents a fundamental challenge in marine microbiology and naturally transcends the species-level heterogeneity of the prokaryotic ocean. Because core metabolic pathways are conserved across ecologically and phylogenetically diverse lineages, the functional signals captured by the KOPAs protocol are transferable across biogeographic provinces, from oligotrophic gyres to productive upwelling systems, making the protocol inherently global in scope. Analysis of KO abundance patterns resolves the spatio-temporal dynamics of ecosystem functioning, uncovering the molecular machinery that governs marine biogeochemical processes. As prokaryotic communities respond rapidly to environmental perturbations, these molecular proxies have the potential to serve as early warning indicators of ecosystem regime shifts, detecting changes in ecosystem services distribution.

## Discussion

Collectively, our results establish a bridge between Earth observation and the abundance of microbial functions. By linking information-rich but spatially sparse data, such as metagenomics data, with global satellite observations, KOPAs protocol enables the prediction of molecular function abundances and associated ecosystem services at the ocean surface. This framework provides a scalable tool for monitoring marine productivity, carbon fluxes, and functional diversity at global scales, from space-derived ecological indicators. In addition, this pipeline allows real-time observation of the projected distribution of key plankton-derived indicators based on molecular processes, opening new opportunities for ocean protection. This tool fills a major gap in marine ecosystem conservation and comes at the right time to align with the political will to protect ecosystem services at a global scale. It further offers a better understanding of the biological machinery fueling NPP, carbon export, and functional diversity.

The KOPAs protocol links ecosystem indicators to distinct molecular processes: photosynthetic and respiratory pathways underpin NPP, amino acid and nucleotide metabolism reflects functional diversity, and membrane transport, secretion and signalling pathways are associated with carbon export. We further show that a subset of these molecular functions can be inferred from satellite observations and projected globally, allowing the spatio-temporal dynamics of ecosystem functioning to be reconstructed while retaining a biological link to the processes that underpin it. Integrating these indicators reveals key oceanic regions where high NPP, carbon export, and functional diversity converge, highlighting locations where multiple plankton-derived ecosystem services are simultaneously supported. Their spatial patterns are consistent with established oceanographic processes, including mesoscale frontal activity and coastal upwelling, while also revealing less-studied regions where the concurrence of these biological signals warrants further investigation.

Despite the capacity of the developed KOPAs protocol to robustly capture the spatial structure of ecosystem services such as NPP and major ecological gradients, their projections remain relative: uncertainty is associated with the magnitude of predictions (Figure 2A and 3C). This limitation is not unique to the KOPAs protocol; quantitative NPP estimates derived from satellite data are inherently uncertain, due to both heterogeneity in satellites observation and variability in model complexity, structure, and ecological assumptions, as described in Supplementary material 10. Another noteworthy limitation comes from the availability of training data. In this study, the limited availability of training data for the carbon export models resulted in an *R*^2^ of approximately 0.2 (Figure 2A) and overfitting across folds in KO to satellite and KO to indicator predictions. These high levels of overfitting are causing low reliability in generalizing carbon export predictions. This highlights the need for tighter coupling between carbon export measurements and omics data collection, and coordinated sampling campaigns that target export flux alongside microbial functional profiling, particularly in regions characterized by extreme environmental conditions, where models are likely to perform poorly in the absence of reference data.

Additionally, integrating multiple omics data sources introduces inherent variability in sequencing depth across oceanographic missions. Therefore, the detection of rare KOs, and thus Shannon diversity estimates, varies among cruises (average Shannon index values ranged from 7.37 for GEOTRACES to 7.56 for Malaspina, see Figure S6). However, as KO abundances are weighted by the relative abundances of their associated MAGs, the influence of rare KOs on the overall indicators is substantially minimized, mitigating this potential bias.

Complementary uncertainties arise from the satellite data themselves [46]. Indeed, MODIS ocean color observations suffer from significant spatial and temporal sampling limitations due to cloud cover, zenith angles and other observing conditions, which introduce biases in chlorophyll a estimates. These biases are particularly severe at high latitudes, primarily due to the exclusion of data at solar zenith angles *>*75° in the processing algorithms.

### Perspectives for marine governance

To assess the robustness of the current governance framework, we examined the interaction between KOPAs’ coverage defined through our protocol and current marine protected areas (MPAs) coverage. We notice that the percentage of high functional diversity values within MPAs exceeds the percentage of MPAs coverage, indicating that the current MPAs distribution better captures plankton-based functional diversity than would be expected if MPAs were randomly placed (Figure 5B). This could be explained by the environmental heterogeneity of coastal ecosystems, where MPAs are predominantly located. We observed, however, that the current MPAs design yields lower NPP and carbon export values than a randomly placed design would. We noticed a change across months in the capacity of MPAs to capture KOPAs (Figure 5C). In addition, our results highlight the variability across countries, with the annual average proportion of MPAs area covered by at least one indicator among NPP and functional diversity, ranging from 18.2% to 90.1% (Figure 5D and S8). For instance, on average, 75% of high levels of NPP and functional diversity occurring in Uruguay’s economic exclusive zones (EEZ) are located in MPAs, but with a temporal variability ranging from 0% to 100% across months (Figure 5D).

The results presented here have direct implications for the design, review, and implementation of Area-Based Management Tools (ABMTs) in marine conservation [47]. It demonstrates the opportunities for some countries to explore the placement of their future MPAs in a way that maximizes coverage of the high projected indicator values happening within their EEZ. Beyond future planning, the comparison between the KOPAs defined here and existing MPAs reveals a measurable mismatch between current conservation designations and the distribution of functionally significant planktonic processes. Areas simultaneously exhibiting high values across the three indicators remain largely outside existing protected areas, with inclusion rates ranging from 3.5% to 9.5% depending on the month considered (Figure S7). This result provides a quantitative basis for assessing the adequacy of existing conservation measures and for identifying areas whose ecological importance may not be captured by species-, habitat-, or biodiversity-based criteria alone (Figures S7, S8 and S9). Such information is directly relevant to the review mechanisms established under Article 26 of the BBNJ Agreement and to the monitoring requirements associated with the Kunming–Montreal Global Biodiversity Framework. Importantly, from a governance perspective, the protocol developed in this work generates these distributions through a reproducible evidentiary chain linking molecular functional capacities (KOs), ecological indicators, and satellite-derived environmental observations. This feature is particularly significant from a governance perspective. Conservation decisions are frequently constrained by the limited spatial and temporal coverage of *in situ* observations, especially in ABNJ. The KOPAs protocol provides a standardized methodology capable of generating recurrent assessments at global scales while maintaining a direct connection to biologically meaningful processes. As a result, indicators associated with biological productivity, carbon export, and functional diversity can be monitored continuously through a framework that is both scientifically transparent and operationally scalable.

Taken together, these findings suggest that process-based ecological evidence can now be incorporated into conservation decisions at the spatial and temporal scales at which marine governance operates. The principal implication is not the replacement of existing conservation categories, but the expansion of the evidentiary basis upon which they are applied. Through the integration of metagenomic functions, ecological indicators, and Earth observations, KOPAs protocol provides a framework for identifying, reviewing, and adapting conservation measures based on observed distribution of ecologically significant processes. In this respect, the framework offers a practical pathway for integrating functional ecological information into the implementation of ABMTs under the BBNJ Agreement and into future assessments of progress toward global marine conservation targets.

## Methods

A total of 1711 samples from all main marine biomes and originating from major oceanographic campaigns (TARA Oceans, TARA Polar Circle, Malaspina, Bio-GOSHIP, and GeoTRACES, see Figure S1) were retrieved from the ENA repositories. The metagenomic profiling of these data was performed using the k-mers-based tools Sylph (Shaw and Yu, 2025), and the marine prokaryotic Metagenome Assembled Genomes (MAG) catalog OMDB (available at https://omdb.microbiomics.io/; Paoli et al., 2022), dereplicated at 99% Average Nucleotide Identity (ANI). The resulting MAG catalog comprised 84,670 MAGs at the time of the analyses. The resulting table contained semi-quantitative MAG distribution at the global scale. The final table was further filtered to only keep High Quality MAGs (completeness *>* 90% and contamination*<* 5%). By combining this table with the functional annotation of the MAGs (obtained using the eggnog mapper), a second table containing the global distribution of the KEGG Orthology (KO) terms at global scale was generated, to allow for a functional mapping of the oceans that preserved its links to taxonomic information. To further maintain the semi-quantitative nature of the data, the value for a given KO in a sample was set to the sum of the relative abundances of all MAGs carrying this KO in that sample. If we assume n MAGs are carrying a KO*_p_* in a sample *S*, then:

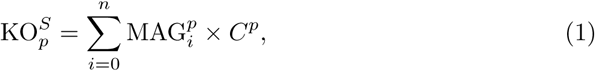

where KO*^S^_p_* is the value for KO*_p_* in sample *S*, MAG*^p^* is the relative abundance of a MAG carrying the KO*_p_*, and *C^p^* is the number of copies of that KO in MAG*^p^*. As such, information about the relative abundances of KOs in each sample was kept, thus mitigating the potential over-representation of KOs found in many extremely rare MAGs compared to KOs found in abundant MAGs.

Consistently with the depth coverage of satellite imagery data, only surface layer samples (0-10 m) were processed (Figure S1) to assess the predictive power of marine microbial functional profiles for key oceanographic parameters. KOs were further filtered so that only individual KOs with a variability *>* 5th quantile were selected. We applied a count-zero multiplicative method to the remaining KOs [1] before performing a CLR transformation on relative abundances as standardized functional features.

### Ecological indicators

Three ecological indicator values (Carbon export, NPP, and functional diversity) were estimated at the omics sampling station when possible. The relative abundance of KOs was used to calculate functional diversity using a Shannon index, reflecting both the richness and evenness of molecular functions within a plankton community (Eq. 2, [2]):

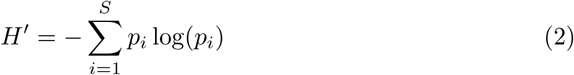

where *p_i_*is the relative abundance of KO *i* and *S* is the total number of KOs observed in the sample.

Concurrent satellite-derived NPP data from the VGPM model [3], https://orca.science.oregonstate.edu, were matched spatio-temporally and extracted from weekly projections at 1080 x 2160 grid resolution for each sampling station. Estimates of particulate carbon export at 150 meters depth were derived from particle concentrations and size distributions measured by the Underwater Vision Profiler [4], they were sourced from *in-situ* measurements and matched to the nearest omics sample with a maximal spatial distance of 7.5km and an identical sampling date.

#### Remote sensing observation

Satellite-derived Level 3 and 4 environmental variables were extracted for each *in-situ* surface sample at its date and geographic location. To obtain environmental data for as many microbial sampling events as possible, we acquired satellite-derived parameters from the NASA OceanColor database (AQUA-MODIS sensor, 4km resolution, 32-day rolling-average composites, recalculated every 8 days). The satellite data included particulate backscattering at 443nm (bbp443), photosynthetically available radiation, day-averaged (PAR), total photosynthetically available radiation (iPAR), sea surface temperature (SST), Chlorophyll a, and the diffuse attenuation coefficient for down-welling irradiance at 490 nm (kd490). For each *in-situ* sample date, we identified the composite files whose temporal coverage window included that date. We selected the file with the period end date closest to the sampling date to maximize temporal relevance. Chlorophyll-a served as the primary proxy for phytoplankton biomass, while bbp443 further discriminated the contribution of non-algal particles (heterotrophic bacteria and organic detritus) to the optical signal. Light availability was assessed through PAR (daily-averaged surface flux) and its vertically-integrated counterpart iPAR, which accounts for in-water attenuation to better represent the total radiant energy accessible to the entire photic-zone community; this was complemented by kd490, an indicator of water clarity that directly constrains the depth of the euphotic layer. Finally, SST was included as a variable controlling metabolic processes and stratification strength.

### From omics to ecosystem indicators prediction

#### KOs selection

To identify KO groups within the surface ocean microbiome, we performed a Weighted Gene Co-expression Network Analysis (WGCNA) on CLR-transformed observed KO abundances. We applied a similarity metric to construct a signed co-expression network based on *ρ* proportionality metric (Eq. 3) to calculate the adjacency matrix. We used a proportionality metric rather than a correlation metric to avoid spurious associations that can arise when correlations are applied to compositional data [5, 6]. The metric *ρ* accounts for the logarithmic relationship among species relative [7, 8].

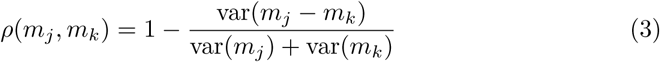

A soft power threshold was selected to approximate a scale-free topology. The threshold value of 5 was applied to the similarity matrix to calculate a Topological Overlap Matrix (TOM). Hierarchical clustering of the TOM-based dissimilarity matrix, followed by dynamic tree cutting, was used to define distinct gene modules, each representing a cluster of highly co-expressed genes. Modules with highly correlated eigengenes were subsequently merged to finalize the network structure.

We related the identified gene modules to NPP, carbon export and functional diversity by calculating Pearson correlations between each module’s eigengene and measured values of the indicators. The statistical significance of module-trait relationships was assessed using Student’s t-test, with p-values adjusted for multiple comparisons using the Benjamini-Hochberg false discovery rate (FDR) correction. Each KO included in a module with a correlation *>* 0.3 with a p-value *<* 0.05 would pass this step of the filtering process.

A second filtration step was applied to the NPP and functional diversity subsets of KOs to identify those most strongly associated with the indicators. We applied Partial Least Squares Regression (PLSr) on the subset of KOs selected by the WGCNA. The optimal number of latent components was determined using a 5-fold cross-validation, minimizing the root mean squared error (RMSE) while monitoring the coefficient of determination (*R*^2^). Model performance was evaluated using both *R*^2^ and RMSE on held-out validation sets. Variable Importance in Projection (VIP) scores were computed for each KO, with VIP *>* 1 used as the threshold to identify genes that contribute significantly to the predictive model.

Extraction of KEGG pathways for all 2589 unique KOs was performed with the KEGGREST package v1.46.0 [9] in R v4.4.1 [10]. Pathway hierarchy was obtained from https://www.genome.jp/kegg/pathway.html. Pathway dereplication and plotting were achieved following the script deposited on the project’s Gitlab. All relevant data regarding functional characterization of KOs associated to ecological indicators can be found on the project’s Gitlab. Pie charts were plotted with ggplot2 v3.5.2 [11].

#### Machine learning predictions

For a KO to be usable in the KOPAs protocol, its abundance must be predicted using remote sensing data. To identify those that match this condition, we employed a nested cross-validation strategy to optimize and evaluate the multi-output XGBoost model [33]. Only samples with an indicator value were used as inputs for the model. In the inner loop, a 3-fold grid search was performed to tune key hyperparameters across a defined parameter space. Due to the limited data on carbon export, a simplified grid search was used. The grid search was configured to maximize the coefficient of determination (*R*^2^) for each outer training fold, independently selecting the optimal hyperparameter set for each of the five outer splits. The best hyperparameter combination was determined by the mode among all splits. This approach ensured that hyperparameter tuning was performed without data leakage from the held-out validation sets. Generalization performance was assessed using an outer 5-fold cross-validation. For each outer fold, the model was trained on the corresponding training subset using the optimal hyperparameters identified in the inner loop, and predictions were generated for the unseen test set. Model performance was evaluated at the individual KO level across all outer folds, aggregating per-KO metrics including *R*^2^ and RMSE. Only the KOs with *R*^2^ *>* 0.3 for NPP and functional diversity, and an *R*^2^ *>* 0.2 for carbon export were selected for further analysis. Additionally, we computed an overfitting gap for each gene, defined as the difference between the average training *R*^2^ and test *R*^2^ across folds. This metric provided a quantitative measure of model stability.

Once the subsets of KOs were identified as relevant to the ecosystem indicator and predictable using remote sensing data, we estimated each subset’s ability to predict the associated indicator. For this purpose, we trained an XGBoost regression model using a repeated cross-validation strategy. Hyperparameter tuning was performed using 5-fold cross-validation, repeated 5 times. Once the optimal hyperparameters were determined, we performed repeated 5-fold cross-validation on observed KO abundances to predict the log(1p)-transformed indicator, thereby increasing the stability of projections. The model was regularized and included early stopping (50 rounds for functional diversity, 25 for NPP and 10 for carbon export) based on validation RMSE. Performance was assessed using the coefficient of determination (*R*^2^) and RMSE. Overfitting was monitored as the difference between training and validation *R*^2^; a gap exceeding 0.2 was flagged as overfitting. This pipeline allowed us to quantify how well a compact set of functionally relevant genes could predict a key ecological indicator, thereby linking genomic potential to biodiversity levels and ecosystem functions. In addition, we performed a Spearman’s rank correlation on carbon export output to assesses the strength and direction of monotonic associations between two observed values and KOPAs outputs. A final step involved projecting the respective KO subsets at the global scale and deriving the associated indicator value. Projections were made on a rolling 32-day average, with a spatial resolution of 4km.

#### Marine Protected Areas

The projections of NPP, carbon export, and functional diversity were compared with MPAs designs to assess the current MPAs pattern’s ability to capture plankton-based ecosystem services and functional diversity. We used MPAtlas from the Marine Conservation Institute because it focuses specifically on MPAs that are both fully protected from extraction and actively implemented at sea (MPAtlas.org). For each indicator, we selected values above the 70th quantile, keeping only the highest values for each parameter. Then we conducted a comparative analysis of the high-value indicator level distributions by computing the proportion of high indicators located in MPAs and comparing it with the proportion of the surface covered by MPAs.

#### Data and code availability

All code required to reproduce the analyses and figures presented in this study is publicly available at https://gitlab.univ-nantes.fr/serandour-b/kopas_serandour/, together with the computational environment used for the analyses. The underlying satellite datasets are publicly available from NASA Ocean Color website (https://oceandata.sci.gsfc.nasa.gov/l3/) and NPP values are publicly available on: https://orca.science.oregonstate.edu/npp_products.php. Biological datasets including metagenomics can be found here: [repository/Zenodo DOI].

## Authors contribution

D.E., A.A., A.M. and B.S. conceptualised the project. B.S. developed the methodology.

B.S. performed the formal analyses and visualisation. D.E. contributed to the develpment and refinement of the methodology and interpretation of the results. C.M.A-G. analysed the functional roles of KOs. A.B. collected metagenomic data from publicly available databases and curated the resulting datasets. R.EH. provided advice on the use of satellite data. S.B-G provides legal implications of KOPAs. B.S. wrote the original draft. A.A. secured funding. A.C., L.G., I.H. and S.C. provided constructive comments and contributed to the review and editing of the manuscript. All authors contributed to review and edit the manuscript.

### Acknowledgments

The authors are also grateful to Alessandro Tagliabue for his valuable comments on the selection and interpretation of net primary production (NPP) sources. Computational support was provided by the bioinformatics core facility of Nantes (BiRD-Biogenouest), Nantes Université, France.

## Supplementary material

**Fig. S1.**
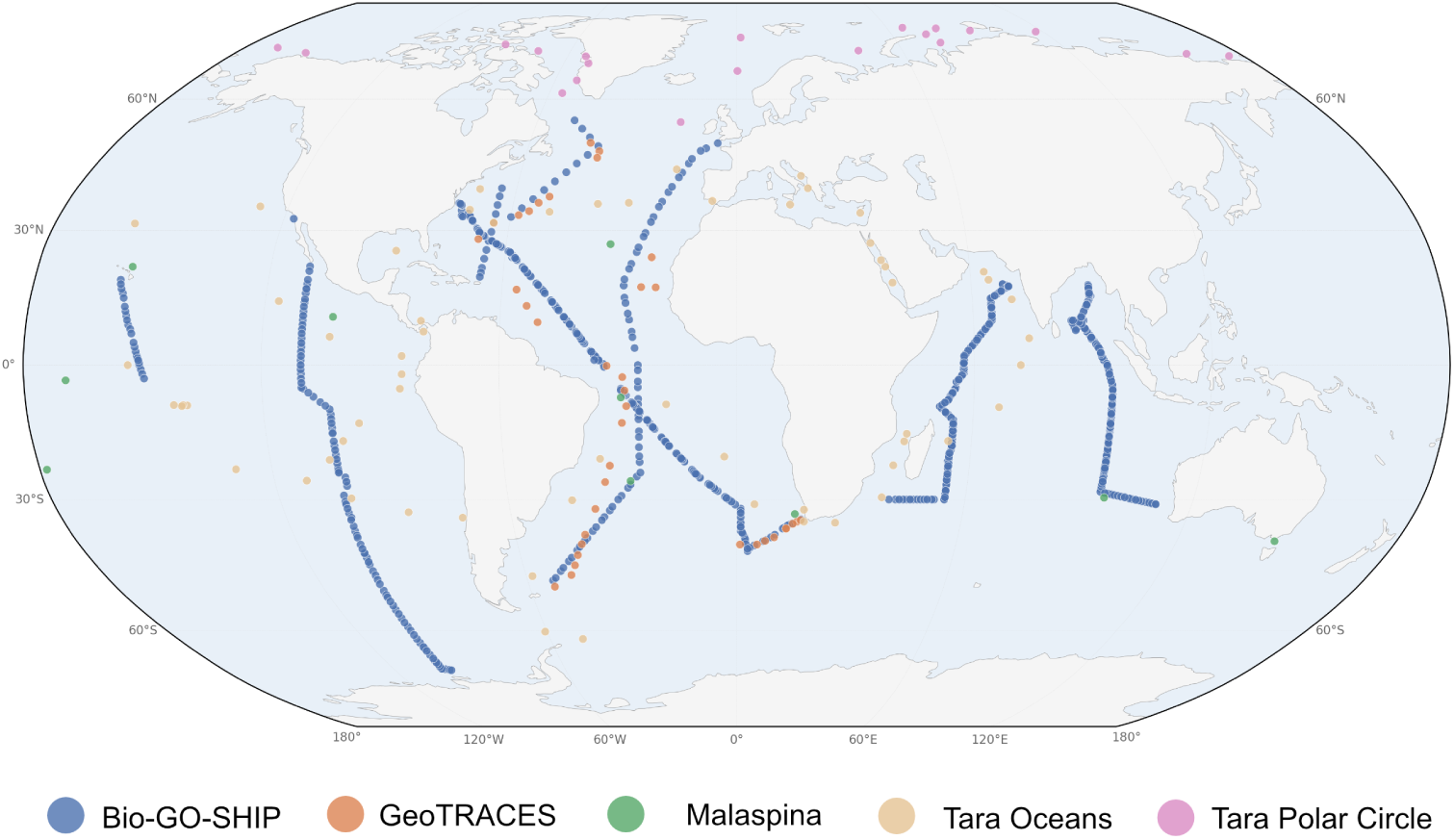
Sampling map. Samples from five major oceanographic campaigns (TARA Oceans [1], TARA Polar Circle [2], Malaspina [3], Bio-GO-SHIP [4], and GeoTRACES [5] have been combined to cover all oceans and biomes.

**Fig. S2.**
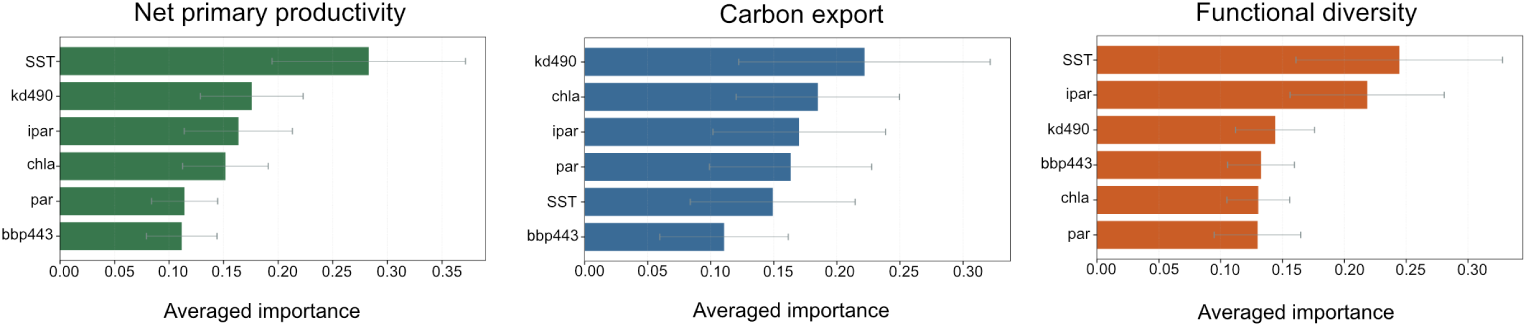
Variable importance for KOs predictability using satellites data. Averaged variable importance for the projections of each subsets of KOs per indicator.

**Fig. S3.**
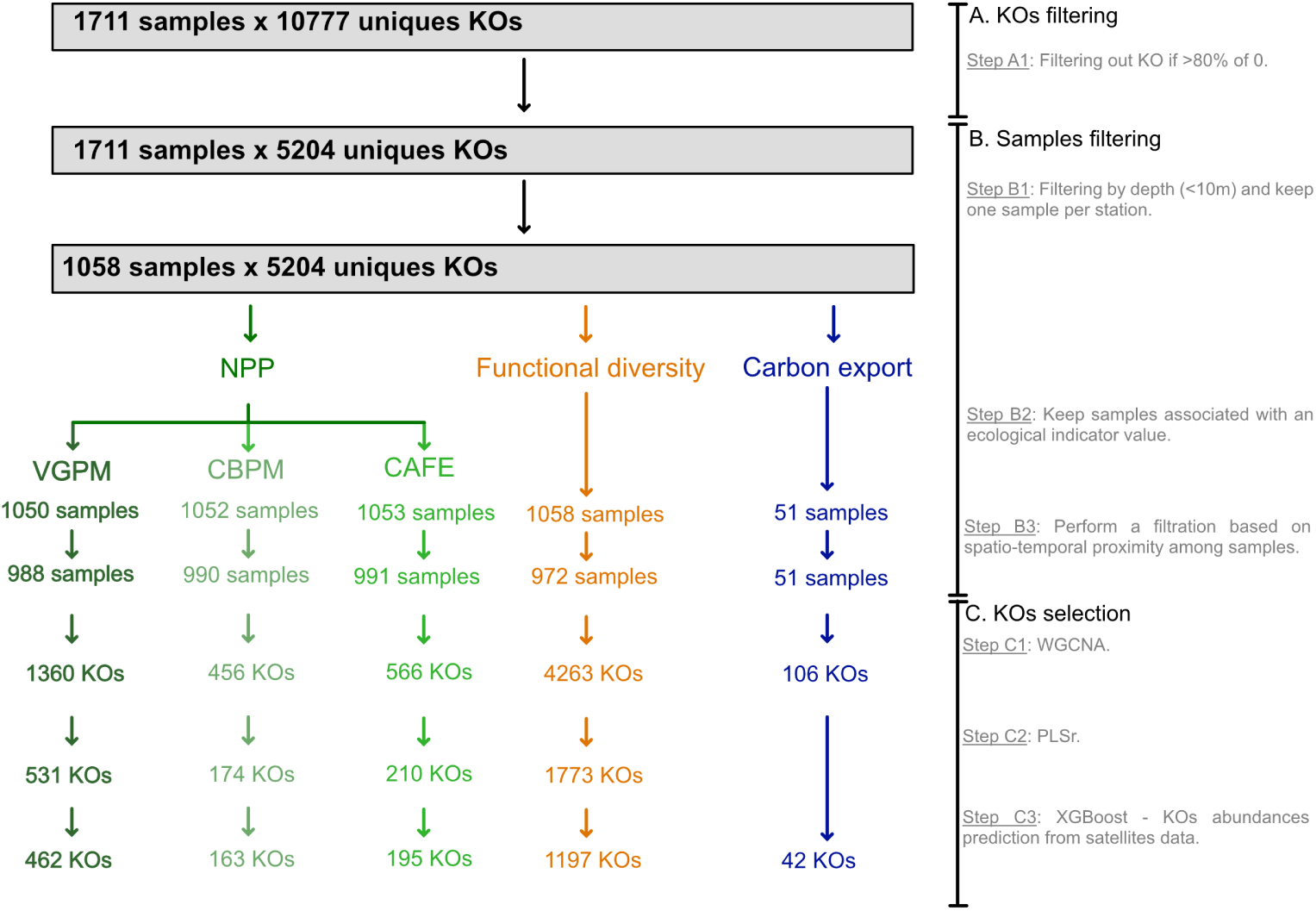
Samples and KOs filtration steps in the KOPAs protocol. Progressive filtering of samples and KOs throughout the KOPAs analytical pipeline, showing the number of samples and KOs retained at each filtering step.

**Fig. S4.**
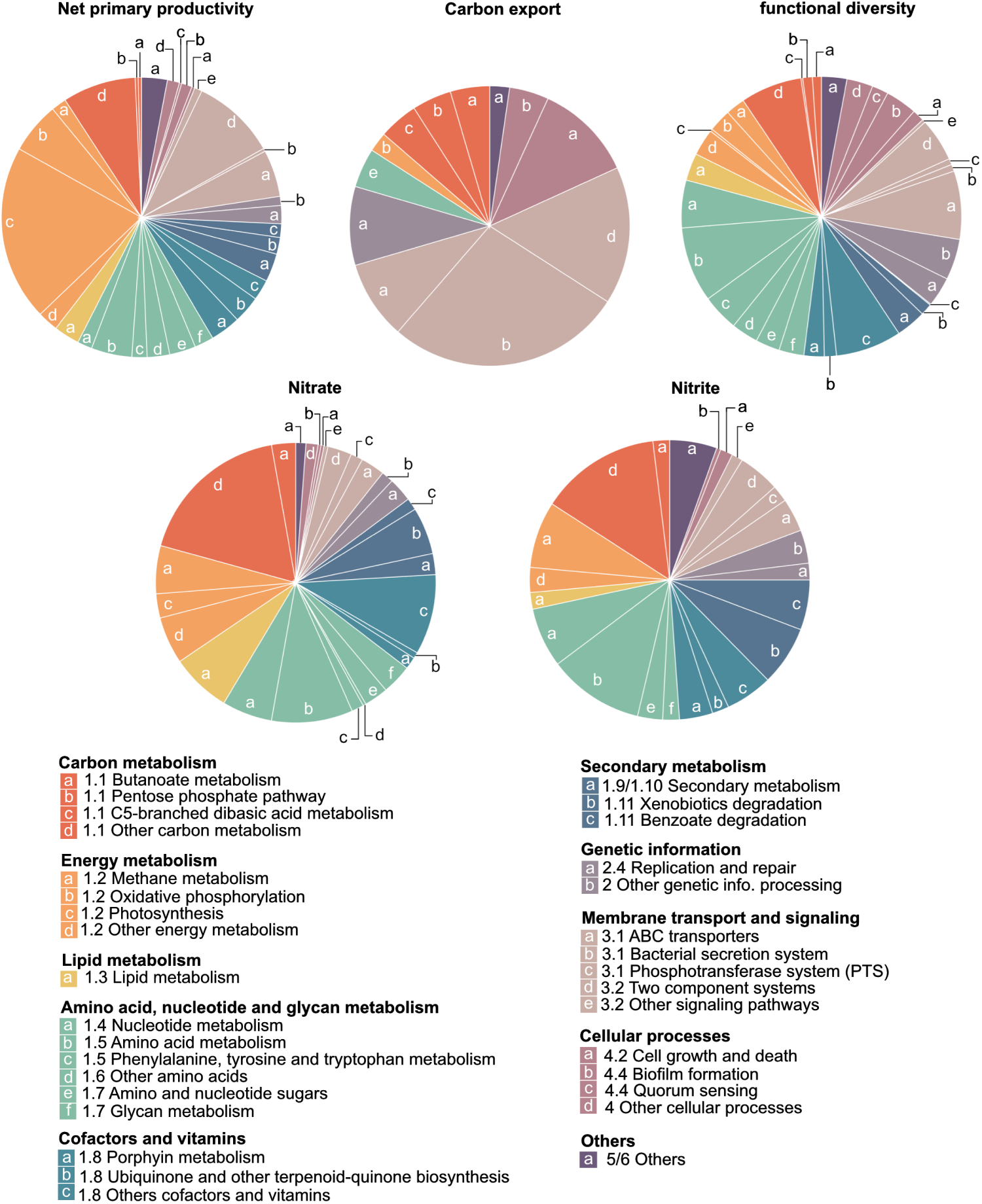
KEGG pathways contained in KOs subsets. Projection of selected KOs onto pathway maps for net primary production, carbon export, and functional diversity. Nitrate and nitrite pathways were included as internal benchmarks to assess the biological coherence and statistical robustness of the selection process. Their well-characterised ecological roles enabled verification that the combined WGCNA–PLSr approach (VIP 1.0) correctly recovers known functional signatures, thereby reinforcing confidence in the less-characterised indicator pathways. Note that pathways were either kept, collapsed, or dropped for improved visualization (see Table kos to pathways on the project’s GitLab).

**Fig. S5.**
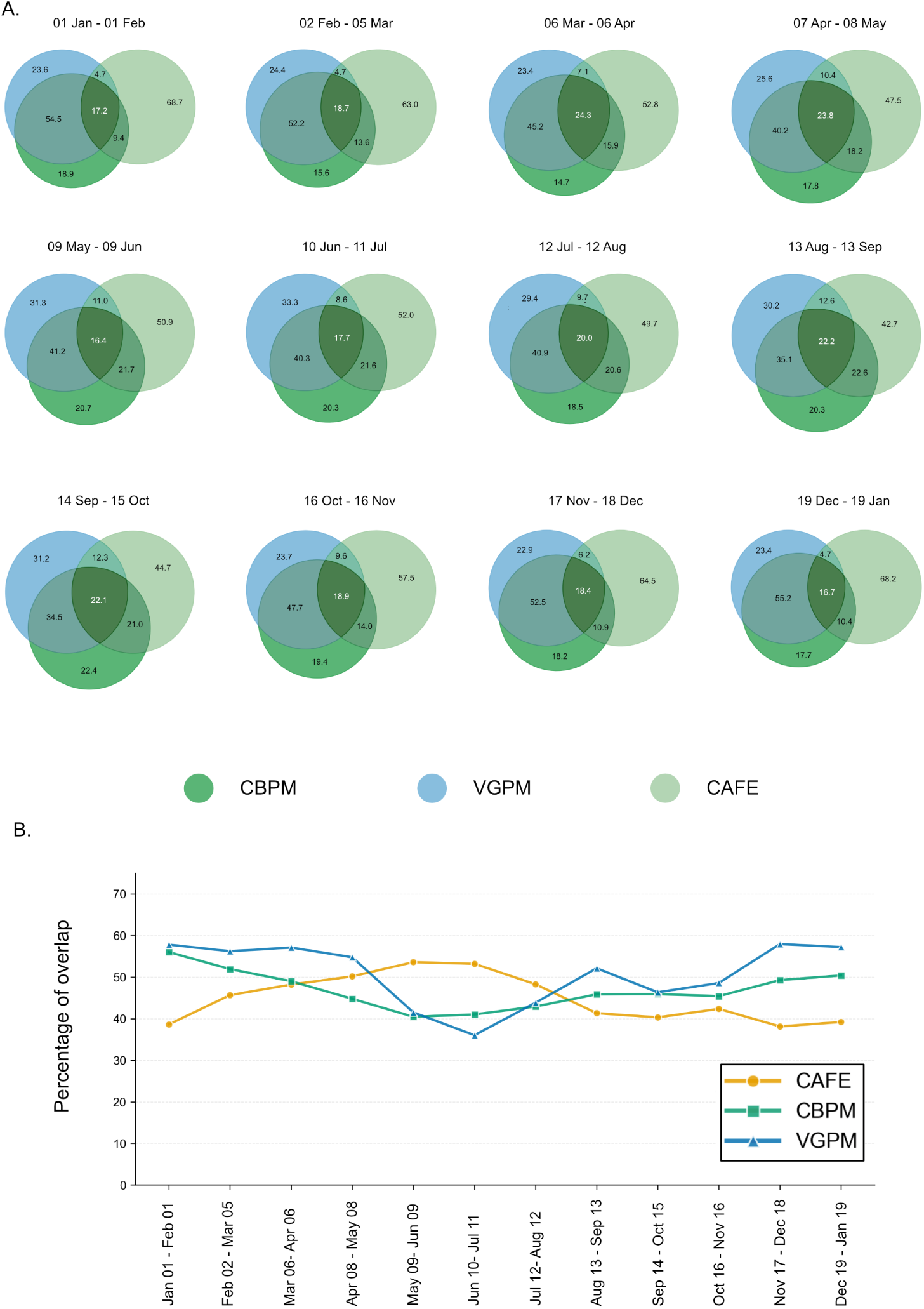
Comparison of NPP high indicator surface across NPP models. **a** Proportion of overlap of high indicator KOPAs-based surface across NPP projections. **b** Proportion of overlap between KOPAs-based projections high indicator, using VGPM [6], CAFE [7] and CBPM [8] and their the high indicator from the original models.

**Fig. S6.**
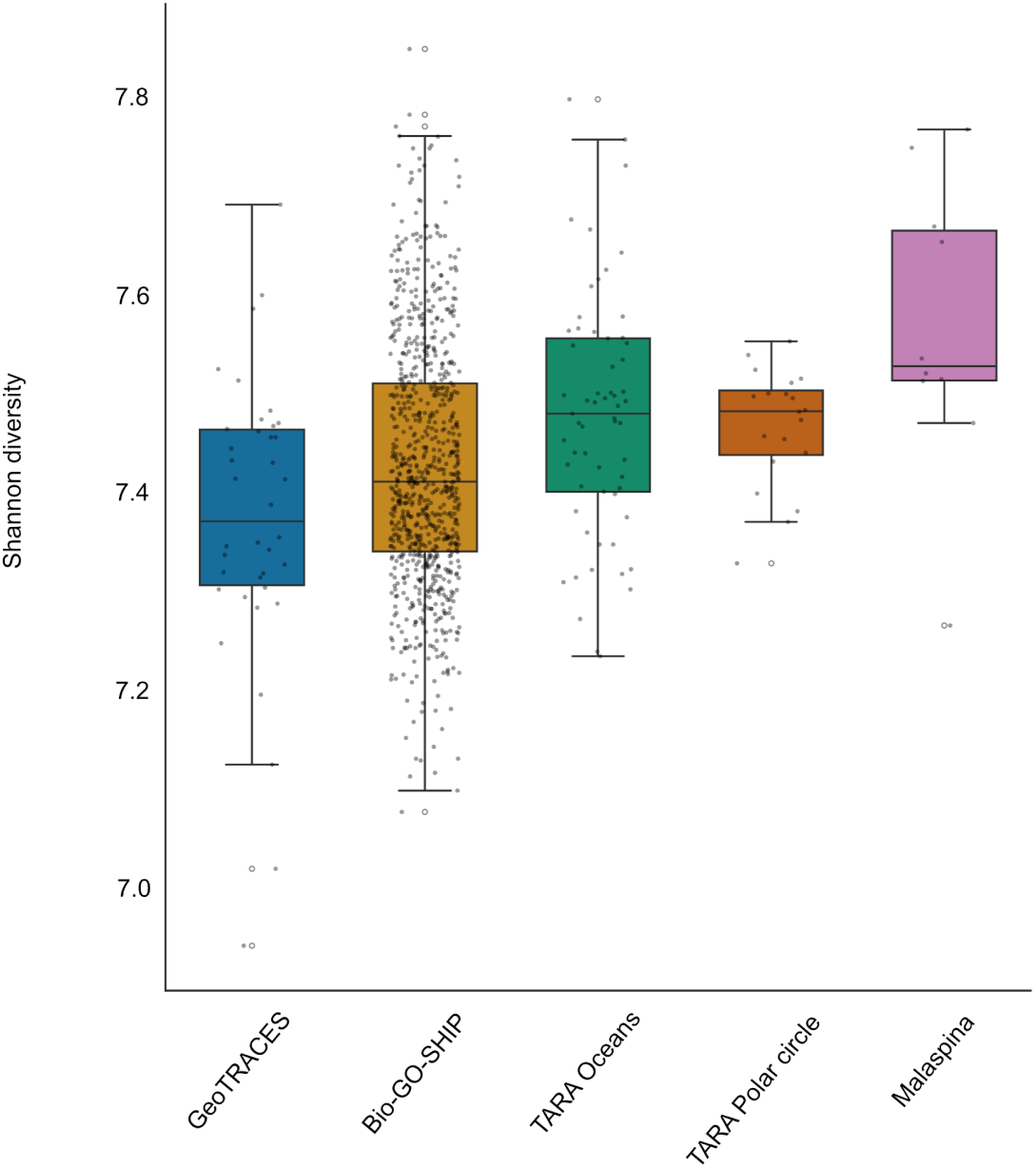
Comparison of Shannon diversity values across oceanographic missions. Variations in sequencing depth across missions may affect the detection of low-abundance KOs. However, the observed differences in Shannon diversity also reflect genuine ecological variability. For example, despite similar sampling and sequencing protocols, the Tara Oceans, Tara Polar Circle and Malaspina expeditions yielded markedly different average Shannon diversity values. A Kruskall-Wallis with Dunn post hoc statistic test shows statistical differences (p*<*0.05) for Bio-GO-SHIP with Malaspina and Tara oceans and for GeoTRACES with Malaspina and Tara oceans

**Fig. S7.**
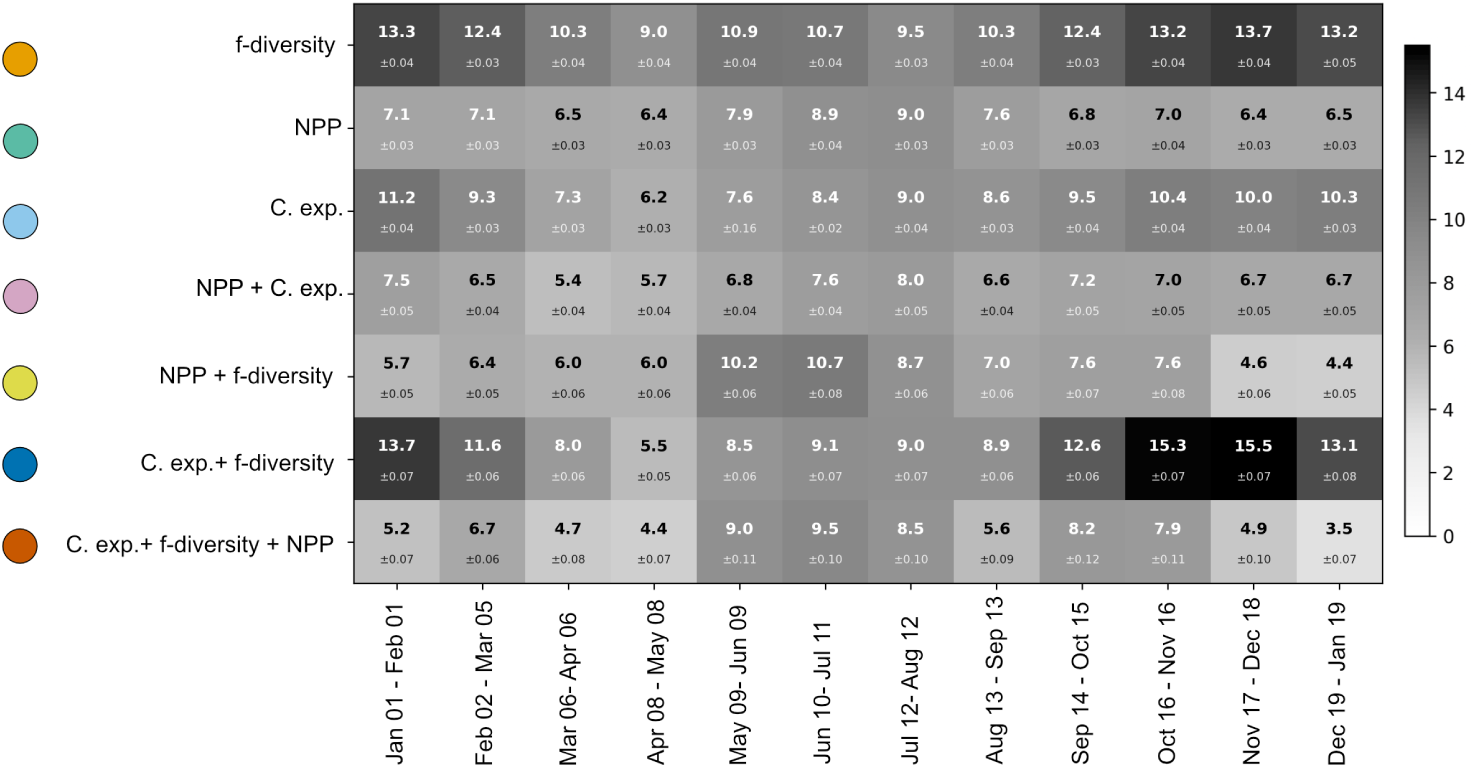
Surface proportion of indicator combination included in MPAs. Indicator combinations are not mutually exclusive: pixels belonging to a multi-indicator combination are also included in each constituent single-indicator set. Notably, the proportion of protected surface within a given combination is not constrained to decrease with the number of indicators, as the spatial co-occurrence of high-value pixels across multiple indicators may preferentially overlap with existing MPAs, thereby yielding a higher protection rate despite a smaller total surface. A random subsampling was of 2,000,000 points was used with 60 different seeds.

**Fig. S8.**
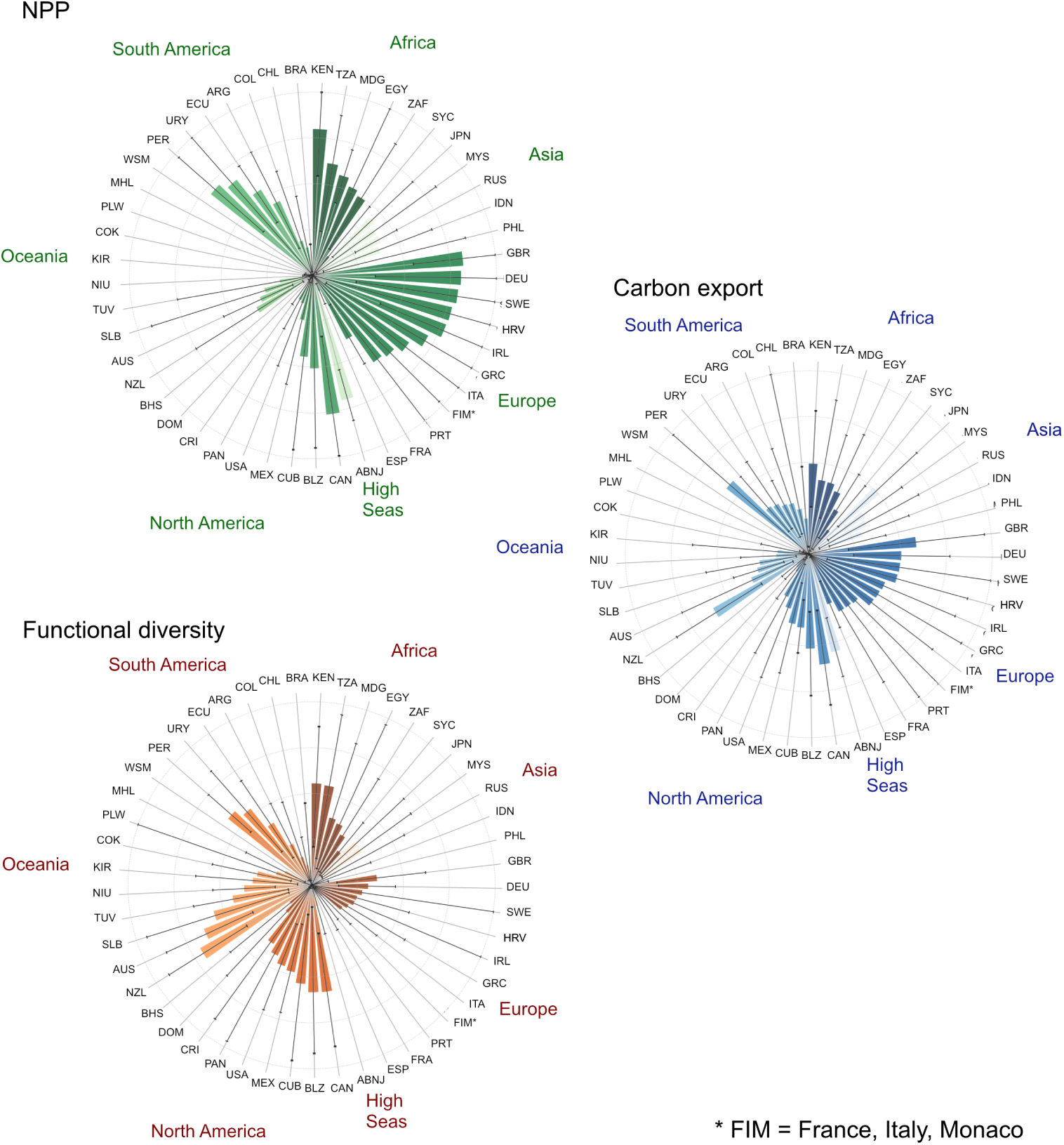
Averaged surface proportion of MPAs of each country covered by high value indicator. Average proportion of MPAs surface covered by high indicator levels with A panel showing NPP, B showing carbon export and functional diversity. The average was calculated on the period from the January 1st 2023 to the January 19th 2024, the error bars show the variability on the period for this country.

**Fig. S9.**
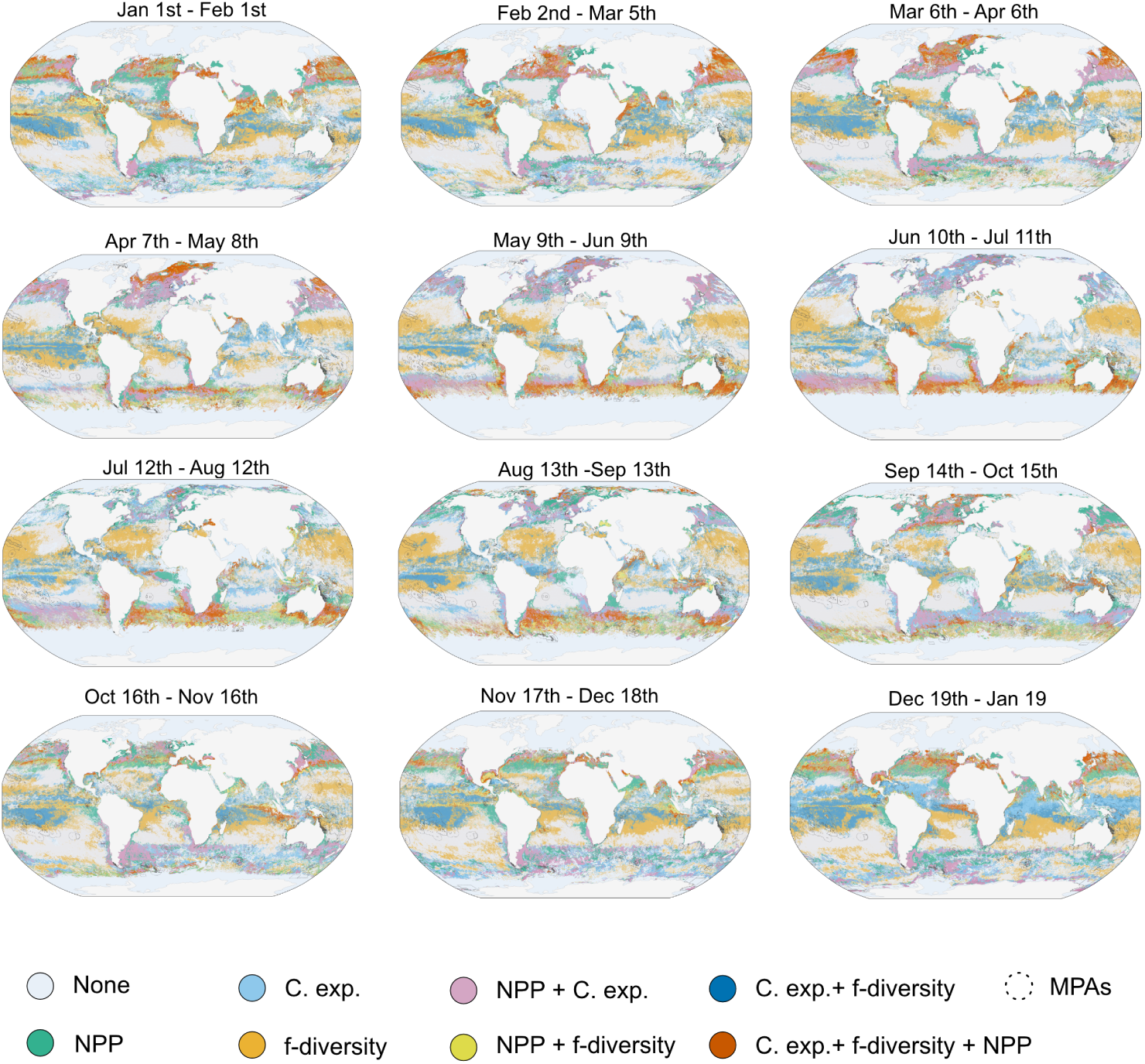
KOPAs distribution across the year 2023. Time series of projected high (*>*70th percentile) NPP, carbon export and functional diversity at 32-day resolution from January 2023 to January 2024.

## Supplementary material 10: Variability in NPP products

In this project, NPP levels computed using the KOPAs protocol, using VGPM as input data, as it remains a standard NPP model in oceanographic studies [9]. However, recent satellite-based NPP models have introduced alternative biophysical formulations, yielding divergent NPP assumptions and estimates [7, 10]. VGPM, CbPM, and CAFE differ fundamentally in how they represent phytoplankton biomass and physiology. VGPM uses surface chlorophyll-a as a proxy for photosynthetically active biomass and parameterizes chlorophyll-normalized photosynthetic rates mainly from sea-surface temperature and light availability. It is therefore sensitive to uncertainties in satellite chlorophyll retrievals, regional variations in the phytoplankton carbon-to-chlorophyll ratio, and the ability of temperature-based parameterizations to represent physiological variability [6]. CbPM instead estimates phytoplankton carbon from particulate backscattering and derives growth rates from chlorophyll-to-carbon variations associated with photoacclimation, light limitation, and nutrient stress. This reduces its reliance on chlorophyll as a biomass proxy but introduces uncertainties related to non-algal contributions to backscattering, the conversion of bbp into phytoplankton carbon, and the physiological interpretation of chlorophyll-to-carbon ratios [10]. CAFE uses an absorption- and energy-based formulation, relating NPP to the amount of photosynthetically available radiation absorbed by phytoplankton and the efficiency with which absorbed energy is converted into carbon. Its estimates are therefore sensitive to the retrieval and partitioning of inherent optical properties, phytoplankton absorption, quantum efficiency, and physiological regulation [7].

The contrasting assumptions among models can produce substantial differences in NPP amplitudes and spatial patterns, even when the models use similar satellite observations. Multi-model comparisons have reported approximately twofold differences in globally integrated production, with particularly strong divergence in cold, high-chlorophyll waters and regions such as the Southern Ocean [11]. Comparisons with *in-situ* NPP have also shown that model performance varies among environmental and optical regimes and that greater spectral or vertical complexity does not necessarily improve predictive skill [12, 13]. Disagreement among products should therefore be interpreted as structural uncertainty arising from alternative representations of phytoplankton biomass, photoacclimation, nutrient limitation, light absorption, and photosynthetic efficiency. Conversely, agreement among products may provide greater confidence in the robustness of large-scale spatial or temporal patterns, even when absolute NPP estimates remain uncertain. Applying the KOPAs framework to three widely used satellite-derived NPP products (VGPM, CbPM and CAFE) confirmed that these methodological differences propagate into conservation prioritization (Figure S5). While the spatial distribution of high-productivity KOPAs was broadly consistent across models, the degree of overlap varied substantially, with VGPM and CbPM showing much greater agreement than either model with CAFE (Fig. S7). This indicates that a fraction of NPP-driven KOPAs reflects structural uncertainty associated with the underlying productivity model rather than ecological variability alone.

## Supplementary material 11: Data leakage in the KOPAs pipeline

Machine-learning algorithms were used at two stages of the KOPAs pipeline: first, to predict KO abundances from satellite-derived observations, and second, to predict ecological indicators from KO abundances. In both cases, model evaluation must be performed on data that are independent of model training and selection; otherwise, information from the evaluation data may enter the modelling process, resulting in data leakage and potentially inflated estimates of predictive performance.

For KO abundance prediction from satellite observations, we addressed this issue using nested cross-validation. In the outer loop, the data were partitioned into folds, with each fold held out once for model evaluation, while the remaining data were used for model development. Within each outer training set, an inner cross-validation procedure was used to select the optimal hyperparameters. This separation between hyperparameter optimisation and final model evaluation provides a reliable and unbiased estimate of predictive performance while reducing the risk of overfitting during model selection. For the prediction of ecological indicators from KO abundances, we used a repeated cross-validation strategy. Repeatedly partitioning the data into training and validation sets provides a more robust estimate of model performance by reducing the dependence of the evaluation on any single data split. Thus, both machine-learning stages were designed to prevent information from the evaluation data from entering model training or hyperparameter selection. However, a potential source of bias remains upstream of the machine-learning steps. WGCNA and PLS regression were performed on the complete dataset prior to the machine-learning procedures. Consequently, KO selection was informed by the ecological indicators before the data were partitioned for model training and evaluation. This constitutes a form of supervised feature selection and may lead to optimistic estimates of downstream predictive performance. The resulting performance estimates should therefore be interpreted in the context of this upstream feature-selection strategy.

## References

[1] Harold Mooney, Anne Larigauderie, Manuel Cesario, Thomas Elmquist, Ove Hoegh-Guldberg, Sandra Lavorel, Georgina M Mace, Margaret Palmer, Robert Scholes, and Tetsukazu Yahara. Biodiversity, climate change, and ecosystem services. Current Opinion in Environmental Sustainability, 1(1):46–54, October 2009. ISSN 18773435. doi: 10.1016/j.cosust.2009.07.006. URL https://linkinghub.elsevier.com/retrieve/pii/S1877343509000086.

[2] R. K. Pachauri, Leo Mayer, and Intergovernmental Panel on Climate Change, editors. Climate change 2014: synthesis report. Intergovernmental Panel on Climate Change, Geneva, Switzerland, 2015. ISBN 978-92-9169-143-2.

[3] María Maestro, Ma Luisa Pérez-Cayeiro, Juan Adolfo Chica-Ruiz, and Harry Reyes. Marine protected areas in the 21st century: Current situation and trends. Ocean & Coastal Management, 171:28–36, April 2019. ISSN 09645691. doi: 10.1016/j.ocecoaman.2019.01.008. URL https://linkinghub.elsevier.com/retrieve/pii/S0964569118305052.

[4] Georgina G. Gurney, Vanessa M. Adams, Jorge G. Aĺvarez Romero, and Joachim Claudet. Area-based conservation: Taking stock and looking ahead. One Earth, 6(2):98–104, February 2023. ISSN 25903322. doi: 10.1016/j.oneear.2023.01.012. URL https://linkinghub.elsevier.com/retrieve/pii/S2590332223000404.

[5] Wu Weiyu. High seas marine protected areas under the bbnj agreement: implementation gaps and governance pathways. Frontiers in Marine Science, 2026. doi: 10.3389/fmars.2026.1860724. URL https://www.frontiersin.org/journals/marine-science/articles/10.3389/fmars.2026.1860724.

[6] Elisabeth Druel and Kristina M. Gjerde. Sustaining marine life beyond boundaries: Options for an implementing agreement for marine biodiversity beyond national jurisdiction under the United Nations Convention on the Law of the Sea. Marine Policy, 49:90–97, November 2014. ISSN 0308597X. doi: 10.1016/j.marpol.2013.11.023. URL https://linkinghub.elsevier.com/retrieve/pii/S0308597X13002820.

[7] Andrew M. Gormley, Elisabeth Slooten, Steve Dawson, Richard J. Barker, Will Rayment, Sam Du Fresne, and Stefan Bräger. First evidence that marine protected areas can work for marine mammals. Journal of Applied Ecology, 49 (2):474–480, April 2012. ISSN 0021-8901, 1365-2664. doi: 10.1111/j.1365-2664.2012.02121.x. URL https://besjournals.onlinelibrary.wiley.com/doi/10.1111/j.1365-2664.2012.02121.x.

[8] K David Hyrenbach, Karin A Forney, and Paul K Dayton. Marine protected areas and ocean basin management. 2000.

[9] Darryn W. Waugh, Edward R. Abraham, and Melissa M. Bowen. Spatial Variations of Stirring in the Surface Ocean: A Case Study of the Tasman Sea. Journal of Physical Oceanography, 36(3):526–542, March 2006. ISSN 1520-0485, 0022-3670. doi: 10.1175/JPO2865.1. URL http://journals.ametsoc.org/doi/10.1175/JPO2865.1.

[10] El Hazen, Rm Suryan, Ja Santora, Sj Bograd, Y Watanuki, and Rp Wilson. Scales and mechanisms of marine hotspot formation. Marine Ecology Progress Series, 487:177–183, July 2013. ISSN 0171-8630, 1616-1599. doi: 10.3354/meps10477. URL http://www.int-res.com/abstracts/meps/v487/p177-183/.

[11] Jeffrey J. Polovina, Evan A. Howell, and Melanie Abecassis. Ocean’s least productive waters are expanding. Geophysical Research Letters, 35(3):2007GL031745, February 2008. ISSN 0094-8276, 1944-8007. doi: 10.1029/2007GL031745. URL https://agupubs.onlinelibrary.wiley.com/doi/10.1029/2007GL031745.

[12] Malin L. Pinsky, Boris Worm, Michael J. Fogarty, Jorge L. Sarmiento, and Simon A. Levin. Marine Taxa Track Local Climate Velocities. Science, 341(6151): 1239–1242, September 2013. ISSN 0036-8075, 1095-9203. doi: 10.1126/science.1239352. URL https://www.science.org/doi/10.1126/science.1239352.

[13] Lionel Guidi, Samuel Chaffron, Lucie Bittner, Damien Eveillard, Abdelhalim Larhlimi, Simon Roux, Youssef Darzi, Stephane Audic, Léo Berline, Jennifer R. Brum, Luis Pedro Coelho, Julio Cesar Ignacio Espinoza, Shruti Malviya, Shinichi Sunagawa, Céline Dimier, Stefanie Kandels-Lewis, Marc Picheral, Julie Poulain, Sarah Searson, Tara Oceans Consortium Coordinators, Lars Stemmann, Fabrice Not, Pascal Hingamp, Sabrina Speich, Mick Follows, Lee Karp-Boss, Emmanuel Boss, Hiroyuki Ogata, Stephane Pesant, Jean Weissenbach, Patrick Wincker, Silvia G. Acinas, Peer Bork, Colomban De Vargas, Daniele Iudicone, Matthew B. Sullivan, Jeroen Raes, Eric Karsenti, Chris Bowler, and Gabriel Gorsky. Plankton networks driving carbon export in the oligotrophic ocean. Nature, 532(7600): 465–470, April 2016. ISSN 0028-0836, 1476-4687. doi: 10.1038/nature16942. URL https://www.nature.com/articles/nature16942.

[14] Robert Ptacnik, Angelo G. Solimini, Tom Andersen, Timo Tamminen, Pål Brettum, Liisa Lepistö, Eva Willén, and Seppo Rekolainen. Diversity predicts stability and resource use efficiency in natural phytoplankton communities. Proceedings of the National Academy of Sciences, 105(13):5134–5138, April 2008. ISSN 0027-8424, 1091-6490. doi: 10.1073/pnas.0708328105. URL https://pnas.org/doi/full/10.1073/pnas.0708328105.

[15] Philip W. Boyd, Hervé Claustre, Marina Levy, David A. Siegel, and Thomas Weber. Multi-faceted particle pumps drive carbon sequestration in the ocean. Nature, 568(7752):327–335, April 2019. ISSN 0028-0836, 1476-4687. doi: 10.1038/s41586-019-1098-2. URL https://www.nature.com/articles/s41586-019-1098-2.

[16] Jerry F. Tjiputra, Damien Couespel, and Richard Sanders. Marine ecosystem role in setting up preindustrial and future climate. Nature Communications, 16 (1):2206, March 2025. ISSN 2041-1723. doi: 10.1038/s41467-025-57371-y. URL https://www.nature.com/articles/s41467-025-57371-y.

[17] D. Pauly and V. Christensen. Primary production required to sustain global fisheries. Nature, 374(6519):255–257, March 1995. ISSN 0028-0836, 1476-4687. doi: 10.1038/374255a0. URL https://www.nature.com/articles/374255a0.

[18] Lydia C L Teh and U R Sumaila. Contribution of marine fisheries to worldwide employment. Fish and Fisheries, 14(1):77–88, March 2013. ISSN 1467-2960, 1467-2979. doi: 10.1111/j.1467-2979.2011.00450.x. URL https://onlinelibrary.wiley.com/doi/10.1111/j.1467-2979.2011.00450.x.

[19] F. Berzaghi, Jérôme Pinti, Olivier Aumont, Olivier Maury, Thomas Cosimano, and Mary S. Wisz. Global distribution, quantification and valuation of the biological carbon pump. Nature Climate Change, 15(4):385–392, April 2025. ISSN 1758-678X, 1758-6798. doi: 10.1038/s41558-025-02295-0. URL https://www.nature.com/articles/s41558-025-02295-0.

[20] Camilo Mora, Derek P. Tittensor, Sina Adl, Alastair G. B. Simpson, and Boris Worm. How Many Species Are There on Earth and in the Ocean? PLoS Biology, 9(8):e1001127, August 2011. ISSN 1545-7885. doi: 10.1371/journal.pbio.1001127. URL https://dx.plos.org/10.1371/journal.pbio.1001127.

[21] Federico M. Ibarbalz, Nicolas Henry, Manoela C. Brandão, Séverine Martini, Greta Busseni, Hannah Byrne, Luis Pedro Coelho, Hisashi Endo, Josep M. Gasol, Ann C. Gregory, Frédéric Mahé, Janaina Rigonato, Marta Royo-Llonch, Guillem Salazar, Isabel Sanz-Sáez, Eleonora Scalco, Dodji Soviadan, Ahmed A. Zayed, Adriana Zingone, Karine Labadie, Joannie Ferland, Claudie Marec, Stefanie Kandels, Marc Picheral, Céline Dimier, Julie Poulain, Sergey Pisarev, Margaux Carmichael, Stéphane Pesant, Silvia G. Acinas, Marcel Babin, Peer Bork, Emmanuel Boss, Chris Bowler, Guy Cochrane, Colomban De Vargas, Mick Follows, Gabriel Gorsky, Nigel Grimsley, Lionel Guidi, Pascal Hingamp, Daniele Iudicone, Olivier Jaillon, Stefanie Kandels, Lee Karp-Boss, Eric Karsenti, Fabrice Not, Hiroyuki Ogata, Stéphane Pesant, Nicole Poulton, Jeroen Raes, Christian Sardet, Sabrina Speich, Lars Stemmann, Matthew B. Sullivan, Shinichi Sunagawa, Patrick Wincker, Marcel Babin, Emmanuel Boss, Daniele Iudicone, Olivier Jaillon, Silvia G. Acinas, Hiroyuki Ogata, Eric Pelletier, Lars Stemmann, Matthew B. Sullivan, Shinichi Sunagawa, Laurent Bopp, Colomban De Vargas, Lee Karp-Boss, Patrick Wincker, Fabien Lombard, Chris Bowler, and Lucie Zinger. Global Trends in Marine Plankton Diversity across Kingdoms of Life. Cell, 179(5):1084–1097.e21, November 2019. ISSN 00928674. doi: 10.1016/j.cell.2019.10.008. URL https://linkinghub.elsevier.com/retrieve/pii/S0092867419311249.

[22] Colleen B. Mouw, Nick J. Hardman-Mountford, Séverine Alvain, Astrid Bracher, Robert J. W. Brewin, Annick Bricaud, Aurea M. Ciotti, Emmanuel Devred, Amane Fujiwara, Takafumi Hirata, Toru Hirawake, Tihomir S. Kostadinov, Shovonlal Roy, and Julia Uitz. A Consumer’s Guide to Satellite Remote Sensing of Multiple Phytoplankton Groups in the Global Ocean. Frontiers in Marine Science, 4, February 2017. ISSN 2296-7745. doi: 10.3389/fmars.2017.00041. URL http://journal.frontiersin.org/article/10.3389/fmars.2017.00041/full.

[23] Astrid Bracher, Heather A. Bouman, Robert J. W. Brewin, Annick Bricaud, Vanda Brotas, Aurea M. Ciotti, Lesley Clementson, Emmanuel Devred, Annalisa Di Cicco, Stephanie Dutkiewicz, Nick J. Hardman-Mountford, Anna E. Hickman, Martin Hieronymi, Takafumi Hirata, Svetlana N. Losa, Colleen B. Mouw, Emanuele Organelli, Dionysios E. Raitsos, Julia Uitz, Meike Vogt, and Aleksandra Wolanin. Obtaining Phytoplankton Diversity from Ocean Color: A Scientific Roadmap for Future Development. Frontiers in Marine Science, 4, March 2017. ISSN 2296-7745. doi: 10.3389/fmars.2017.00055. URL http://journal.frontiersin.org/article/10.3389/fmars.2017.00055/full.

[24] Shovonlal Roy, Shubha Sathyendranath, Heather Bouman, and Trevor Platt. The global distribution of phytoplankton size spectrum and size classes from their light-absorption spectra derived from satellite data. Remote Sensing of Environment, 139:185–197, December 2013. ISSN 00344257. doi: 10.1016/j.rse.2013.08.004. URL https://linkinghub.elsevier.com/retrieve/pii/S0034425713002629.

[25] Hiroto Kaneko, Hisashi Endo, Nicolas Henry, Cédric Berney, Frédéric Mahé, Julie Poulain, Karine Labadie, Odette Beluche, Roy El Hourany, Tara Oceans Coordinators, Silvia G Acinas, Marcel Babin, Peer Bork, Chris Bowler, Guy Cochrane, Colomban De Vargas, Gabriel Gorsky, Lionel Guidi, Nigel Grimsley, Pascal Hingamp, Daniele Iudicone, Olivier Jaillon, Stefanie Kandels, Eric Karsenti, Fabrice Not, Nicole Poulton, Stéphane Pesant, Christian Sardet, Sabrina Speich, Lars Stemmann, Matthew B Sullivan, Shinichi Sunagawa, Samuel Chaffron, Patrick Wincker, Ryosuke Nakamura, Lee Karp-Boss, Emmanuel Boss, Chris Bowler, Colomban De Vargas, Kentaro Tomii, and Hiroyuki Ogata. Predicting global distributions of eukaryotic plankton communities from satellite data. ISME Communications, 3(1):101, December 2023. ISSN 2730-6151. doi: 10.1038/s43705-023-00308-7. URL https://academic.oup.com/ismecommun/article/7584909.

[26] Roy El Hourany, Juan Pierella Karlusich, Lucie Zinger, Hubert Loisel, Marina Levy, and Chris Bowler. Linking satellites to genes with machine learning to estimate phytoplankton community structure from space. Ocean Science, 20(1): 217–239, February 2024. ISSN 1812-0792. doi: 10.5194/os-20-217-2024. URL https://os.copernicus.org/articles/20/217/2024/.

[27] Rebecca Lewison, Alistair J. Hobday, Sara Maxwell, Elliott Hazen, Jason R. Hartog, Daniel C. Dunn, Dana Briscoe, Sabrina Fossette, Catherine E. O’Keefe, Michele Barnes, Melanie Abecassis, Steven Bograd, N. David Bethoney, Helen Bailey, David Wiley, Samantha Andrews, Lucie Hazen, and Larry B. Crowder. Dynamic Ocean Management: Identifying the Critical Ingredients of Dynamic Approaches to Ocean Resource Management. BioScience, 65(5): 486–498, May 2015. ISSN 1525-3244, 0006-3568. doi: 10.1093/biosci/biv018. URL http://academic.oup.com/bioscience/article/65/5/486/323837/Dynamic-Ocean-Management-Identifying-the-Critical.

[28] Sara M. Maxwell, Elliott L. Hazen, Rebecca L. Lewison, Daniel C. Dunn, Helen Bailey, Steven J. Bograd, Dana K. Briscoe, Sabrina Fossette, Alistair J. Hobday, Meredith Bennett, Scott Benson, Margaret R. Caldwell, Daniel P. Costa, Heidi Dewar, Tomo Eguchi, Lucie Hazen, Suzanne Kohin, Tim Sippel, and Larry B. Crowder. Dynamic ocean management: Defining and conceptualizing real-time management of the ocean. Marine Policy, 58:42–50, August 2015. ISSN 0308597X. doi: 10.1016/j.marpol.2015.03.014. URL https://linkinghub.elsevier.com/retrieve/pii/S0308597X15000639.

[29] Morten Frederiksen, Martin Edwards, Anthony J. Richardson, Nicholas C. Halliday, and Sarah Wanless. From plankton to top predators: bottom-up control of a marine food web across four trophic levels. Journal of Animal Ecology, 75(6):1259–1268, November 2006. ISSN 0021-8790, 1365-2656. doi: 10.1111/j.1365-2656.2006.01148.x. URL https://besjournals.onlinelibrary.wiley.com/doi/10.1111/j.1365-2656.2006.01148.x.

[30] Natacha Le Grix and Alessandro Tagliabue. Sources of Uncertainty in Ocean Net Primary Productivity Projections Under Climate Change. Geophysical Research Letters, 53(9):e2025GL119652, May 2026. ISSN 0094-8276, 1944-8007. doi: 10.1029/2025GL119652. URL https://agupubs.onlinelibrary.wiley.com/doi/10.1029/2025GL119652.

[31] P Langfelder and S Horvath. Wgcna: an r package for weighted correlation network analysis. BMC Bioinformatics. doi: 10.1186/1471-2105-9-559. URL https://link.springer.com/article/10.1186/1471-2105-9-559.

[32] Marinna Gaudin, Damien Eveillard, and Samuel Chaffron. Ecological associations distribution modelling of marine plankton at a global scale. Philosophical Transactions of the Royal Society B: Biological Sciences, 379(1909):20230169, September 2024. ISSN 0962-8436, 1471-2970. doi: 10.1098/rstb.2023.0169. URL http://royalsocietypublishing.org/rstb/article/42896.

[33] Tianqi Chen and Carlos Guestrin. Xgboost: A scalable tree boosting system. pages 785–794, 2016. doi: 10.1145/2939672.2939785. URL https://dl.acm.org/doi/10.1145/2939672.2939785.

34. [34] Glen Ellen CA. Marine conservation institute [and partners] mpatlas[on-line] 2026-03-11. 2026. URL mpatlas.org.

[35] Martin F. Hohmann-Marriott and Robert E. Blankenship. Evolution of Photosynthesis. Annual Review of Plant Biology, 62(1):515–548, June 2011. ISSN 1543-5008, 1545-2123. doi: 10.1146/annurev-arplant-042110-103811. URL https://www.annualreviews.org/doi/10.1146/annurev-arplant-042110-103811.

[36] Paul G. Falkowski, Edward A. Laws, Richard T. Barber, and James W. Murray. Phytoplankton and Their Role in Primary, New, and Export Production, pages 99–121. Springer Berlin Heidelberg, Berlin, Heidelberg, 2003. ISBN 978-3-642-55844-3. doi: 10.1007/978-3-642-55844-3_5. URL 10.1007/978-3-642-55844-3_5.

[37] Winifred M. Johnson, Melissa C. Kido Soule, and Elizabeth B. Kujawinski. Evidence for quorum sensing and differential metabolite production by a marine bacterium in response to dmsp. The ISME Journal, 10(9):2304–2316, 2016. ISSN 1751-7370. doi: 10.1038/ismej.2016.6. URL https://www.nature.com/articles/ismej20166.

[38] Xiangyu Wang, Yongxin Lv, Weishu Zhao, Xiang Xiao, and Jing Wang. D-amino acid metabolic versatility as a common adaptive strategy in the mariana trench microbiome. mSystems, 10(8):e00581–25, 2025. doi: 10.1128/msystems.00581-25. URL https://journals.asm.org/doi/abs/10.1128/msystems.00581-25.

[39] Alan W. Decho and Tony Gutierrez. Microbial Extracellular Polymeric Substances (EPSs) in Ocean Systems. Frontiers in Microbiology, 8:922, May 2017. ISSN 1664-302X. doi: 10.3389/fmicb.2017.00922. URL http://journal.frontiersin.org/article/10.3389/fmicb.2017.00922/full.

[40] Carol Arnosti. Microbial Extracellular Enzymes and the Marine Carbon Cycle. Annual Review of Marine Science, 3(1):401–425, January 2011. ISSN 1941-1405, 1941-0611. doi: 10.1146/annurev-marine-120709-142731. URL https://www.annualreviews.org/doi/10.1146/annurev-marine-120709-142731.

[41] Megan Coolahan and Kristen E. Whalen. A review of quorum-sensing and its role in mediating interkingdom interactions in the ocean. Communications Biology, 8(1):179, 2025. ISSN 2399-3642. doi: 10.1038/s42003-025-07608-9. URL 10.1038/s42003-025-07608-9.

[42] Edo Bar-Zeev, Itamar Avishay, Kay D Bidle, and Ilana Berman-Frank. Programmed cell death in the marine cyanobacterium trichodesmium mediates carbon and nitrogen export. The ISME Journal, 7(12):2340–2348, 12 2013. ISSN 1751-7362. doi: 10.1038/ismej.2013.121. URL 10.1038/ismej.2013.121.

[43] Michael J. Behrenfeld and Paul G. Falkowski. Photosynthetic rates derived from satellite-based chlorophyll concentration. Limnology and Oceanography, 42(1):1– 20, January 1997. ISSN 0024-3590, 1939-5590. doi: 10.4319/lo.1997.42.1.0001. URL https://aslopubs.onlinelibrary.wiley.com/doi/10.4319/lo.1997.42.1.0001.

[44] T. Westberry, M. J. Behrenfeld, D. A. Siegel, and E. Boss. Carbon-based primary productivity modeling with vertically resolved photoacclimation. Global Biogeochemical cycles. doi: 10.1029/2007GB003078. URL https://orca.science.oregonstate.edu/references/Westberry_GBC_2008.pdf.

[45] Greg M Silsbe, Michael J Behrenfeld, Kimberly H Halsey, Allen J Milligan, and Toby K Westberry. The cafe model: A net production model for global ocean phytoplankton. Global Biogeochemical cycles. doi: 10.1002/2016GB005521. URL https://orca.science.oregonstate.edu/references/Silsbe_GBC_2016.pdf.

[46] Watson W. Gregg and Nancy W. Casey. Sampling biases in modis and seawifs ocean chlorophyll data. Remote Sensing of Environment, 111:25–35, November 2007. doi: 10.1016/j.rse.2007.03.008. URL https://www.sciencedirect.com/science/article/pii/S0034425707001253.

[47] Elena Gissi, Frank Maes, Zacharoula Kyriazi, Ana Ruiz-Frau, Catarina Frazão Santos, Barbara Neumann, Adriano Quintela, Fátima L. Alves, Simone Borg, Wenting Chen, Maria Da Luz Fernandes, Maria Hadjimichael, Elisabetta Manea, Márcia Marques, Froukje Maria Platjouw, Michelle E. Portman, Lisa P. Sousa, Luca Bolognini, Wesley Flannery, Fabio Grati, Cristina Pita, Natas, a Vaidianu, Robert Stojanov, Jan Van Tatenhove, Fiorenza Micheli, Anna-Katharina Hornidge, and Sebastian Unger. Contributions of marine area-based management tools to the UN sustainable development goals. Journal of Cleaner Production, 330:129910, January 2022. ISSN 09596526. doi: 10.1016/j.jclepro.2021.129910. URL https://linkinghub.elsevier.com/retrieve/pii/S0959652621040804.

## References

[1] Josep Antoni Martin-Fernandez, Karel Hron, Matthias Templ, Peter Filzmoser, and Javier Palarea-Albaladejo. Bayesian-multiplicative treatment of count zeros in compositional data sets. Statistical Modelling, 15(2):134–158, 2015. ISSN 1471-082X. URL https://repositum.tuwien.at/handle/20.500.12708/150426. Accepted: 2023-02-10T07:46:46Z.

[2] C. E. Shannon and W. Weaver. Mathematical theory of communication. 1948.

[3] Michael J. Behrenfeld and Paul G. Falkowski. Photosynthetic rates derived from satellite-based chlorophyll concentration. Limnology and Oceanography, 42(1):1– 20, January 1997. ISSN 0024-3590, 1939-5590. doi: 10.4319/lo.1997.42.1.0001. URL https://aslopubs.onlinelibrary.wiley.com/doi/10.4319/lo.1997.42.1.0001.

[4] Lionel Guidi, Samuel Chaffron, Lucie Bittner, Damien Eveillard, Abdelhalim Larhlimi, Simon Roux, Youssef Darzi, Stephane Audic, Léo Berline, Jennifer R. Brum, Luis Pedro Coelho, Julio Cesar Ignacio Espinoza, Shruti Malviya, Shinichi Sunagawa, Céline Dimier, Stefanie Kandels-Lewis, Marc Picheral, Julie Poulain, Sarah Searson, Tara Oceans Consortium Coordinators, Lars Stemmann, Fabrice Not, Pascal Hingamp, Sabrina Speich, Mick Follows, Lee Karp-Boss, Emmanuel Boss, Hiroyuki Ogata, Stephane Pesant, Jean Weissenbach, Patrick Wincker, Silvia G. Acinas, Peer Bork, Colomban De Vargas, Daniele Iudicone, Matthew B. Sullivan, Jeroen Raes, Eric Karsenti, Chris Bowler, and Gabriel Gorsky. Plankton networks driving carbon export in the oligotrophic ocean. Nature, 532(7600): 465–470, April 2016. ISSN 0028-0836, 1476-4687. doi: 10.1038/nature16942. URL https://www.nature.com/articles/nature16942.

[5] Beiyao Zheng. Summarizing the goodness of fit of generalized linear models for longitudinal data. Statistics in Medicine, 19(10):1265–1275, May 2000. ISSN 0277-6715, 1097-0258. doi: 10.1002/(SICI)1097-0258(20000530)19:10⟨1265::AID-SIM486⟩3.0.CO;2-U. URL https://onlinelibrary.wiley.com/doi/10.1002/(SICI)1097-0258(20000530)19:10⟨1265::AID-SIM486⟩3.0.CO;2-U.

[6] David Lovell, Vera Pawlowsky-Glahn, Juan José Egozcue, Samuel Marguerat, and Jürg Bähler. Proportionality: A Valid Alternative to Correlation for Relative Data. PLOS Computational Biology, 11(3):e1004075, March 2015. ISSN 1553-7358. doi: 10.1371/journal.pcbi.1004075. URL https://dx.plos.org/10.1371/journal.pcbi.1004075.

[7] Ionas Erb and Cedric Notredame. How should we measure proportionality on relative gene expression data? Theory in Biosciences, 135(1-2):21–36, June 2016. ISSN 1431-7613, 1611-7530. doi: 10.1007/s12064-015-0220-8. URL http://link.springer.com/10.1007/s12064-015-0220-8.

[8] Marinna Gaudin, Damien Eveillard, and Samuel Chaffron. Ecological associations distribution modelling of marine plankton at a global scale. Philosophical Transactions of the Royal Society B: Biological Sciences, 379(1909):20230169, September 2024. ISSN 0962-8436, 1471-2970. doi: 10.1098/rstb.2023.0169. URL http://royalsocietypublishing.org/rstb/article/42896.

[9] D Tenenbaum and B Maintainer. Keggrest: Client-side rest access to the kyoto encyclopedia of genes and genomes (kegg). r package version 1.52.2. 2026. doi: 10.18129/B9.bioc.KEGGREST. URL https://bioconductor.org/packages/KEGGREST.

[10] R Core Team. R: A Language and Environment for Statistical Computing. R Foundation for Statistical Computing, Vienna, Austria, 2024. URL https://www.R-project.org/.

[11] Hadley Wickham. ggplot2: Elegant Graphics for Data Analysis. Springer-Verlag New York, 2016. ISBN 978-3-319-24277-4. URL https://ggplot2.tidyverse.org.

## References

[1] Eric Karsenti, Silvia G. Acinas, Peer Bork, Chris Bowler, Colomban De Vargas, Jeroen Raes, Matthew Sullivan, Detlev Arendt, Francesca Benzoni, Jean-Michel Claverie, Mick Follows, Gaby Gorsky, Pascal Hingamp, Daniele Iudicone, Olivier Jaillon, Stefanie Kandels-Lewis, Uros Krzic, Fabrice Not, Hiroyuki Ogata, Stéphane Pesant, Emmanuel Georges Reynaud, Christian Sardet, Michael E. Sieracki, Sabrina Speich, Didier Velayoudon, Jean Weissenbach, Patrick Wincker, and the Tara Oceans Consortium. A Holistic Approach to Marine Eco-Systems Biology. PLoS Biology, 9(10):e1001177, October 2011. ISSN 1545-7885. doi: 10.1371/journal.pbio.1001177. URL https://dx.plos.org/10.1371/journal.pbio.1001177.

[2] Stéphane Pesant, Fabrice Not, Marc Picheral, Stefanie Kandels-Lewis, Noan Le Bescot, Gabriel Gorsky, Daniele Iudicone, Eric Karsenti, Sabrina Speich, Romain Troublé, Céline Dimier, Sarah Searson, Tara Oceans Consortium Coordinators, Silvia G. Acinas, Peer Bork, Emmanuel Boss, Chris Bowler, Colomban De Vargas, Michael Follows, Gabriel Gorsky, Nigel Grimsley, Pascal Hingamp, Daniele Iudicone, Olivier Jaillon, Stefanie Kandels-Lewis, Lee Karp-Boss, Eric Karsenti, Uros Krzic, Fabrice Not, Hiroyuki Ogata, Stéphane Pesant, Jeroen Raes, Emmanuel G. Reynaud, Christian Sardet, Mike Sieracki, Sabrina Speich, Lars Stemmann, Matthew B. Sullivan, Shinichi Sunagawa, Didier Velayoudon, Jean Weissenbach, and Patrick Wincker. Open science resources for the discovery and analysis of Tara Oceans data. Scientific Data, 2(1):150023, May 2015. ISSN 2052-4463. doi: 10.1038/sdata.2015.23. URL https://www.nature.com/articles/sdata201523.

[3] Carlos M. Duarte. Seafaring in the 21St Century: The Malaspina 2010 Circumnavigation Expedition. Limnology and Oceanography Bulletin, 24(1):11–14, February 2015. ISSN 1539-607X, 1539-6088. doi: 10.1002/lob.10008. URL https://aslopubs.onlinelibrary.wiley.com/doi/10.1002/lob.10008.

[4] Sophie Clayton, Harriet Alexander, Jason R. Graff, Nicole J. Poulton, Luke R. Thompson, Heather Benway, Emmanuel Boss, and Adam Martiny. Bio-GOSHIP: The Time Is Right to Establish Global Repeat Sections of Ocean Biology. Frontiers in Marine Science, 8:767443, January 2022. ISSN 2296-7745. doi: 10.3389/fmars.2021.767443. URL https://www.frontiersin.org/articles/10.3389/fmars.2021.767443/full.

[5] Robert F. Anderson. GEOTRACES: Accelerating Research on the Marine Biogeochemical Cycles of Trace Elements and Their Isotopes. Annual Review of Marine Science, 12(1):49–85, January 2020. ISSN 1941-1405, 1941-0611. doi: 10.1146/annurev-marine-010318-095123. URL https://www.annualreviews.org/doi/10.1146/annurev-marine-010318-095123.

[6] Michael J. Behrenfeld and Paul G. Falkowski. Photosynthetic rates derived from satellite-based chlorophyll concentration. Limnology and Oceanography, 42(1):1– 20, January 1997. ISSN 0024-3590, 1939-5590. doi: 10.4319/lo.1997.42.1.0001. URL https://aslopubs.onlinelibrary.wiley.com/doi/10.4319/lo.1997.42.1.0001.

[7] Greg M. Silsbe, Michael J. Behrenfeld, Kimberly H. Halsey, Allen J. Milligan, and Toby K. Westberry. The CAFE model: A net production model for global ocean phytoplankton. Global Biogeochemical Cycles, 30(12):1756–1777, December 2016. ISSN 0886-6236, 1944-9224. doi: 10.1002/2016GB005521. URL https://agupubs.onlinelibrary.wiley.com/doi/10.1002/2016GB005521.

[8] Michael J. Behrenfeld, Emmanuel Boss, David A. Siegel, and Donald M. Shea. Carbon-based ocean productivity and phytoplankton physiology from space. Global Biogeochemical Cycles, 19(1):2004GB002299, March 2005. ISSN 0886-6236, 1944-9224. doi: 10.1029/2004GB002299. URL https://agupubs.onlinelibrary.wiley.com/doi/10.1029/2004GB002299.

[9] Fernando González Taboada, Andrew D. Barton, Charles A. Stock, John Dunne, and Jasmin G. John. Seasonal to interannual predictability of oceanic net primary production inferred from satellite observations. Progress in Oceanography, 170: 28–39, January 2019. ISSN 00796611. doi: 10.1016/j.pocean.2018.10.010. URL https://linkinghub.elsevier.com/retrieve/pii/S0079661117300733.

[10] T. Westberry, M. J. Behrenfeld, D. A. Siegel, and E. Boss. Carbon-based primary productivity modeling with vertically resolved photoacclimation. Global Biogeochemical cycles. doi: 10.1029/2007GB003078. URL https://orca.science.oregonstate.edu/references/Westberry_GBC_2008.pdf.

[11] Mary-Elena Carr, Marjorie A.M. Friedrichs, Marjorie Schmeltz, Maki Noguchi Aita, David Antoine, Kevin R. Arrigo, Ichio Asanuma, Olivier Aumont, Richard Barber, Michael Behrenfeld, Robert Bidigare, Erik T. Buitenhuis, Janet Campbell, Aurea Ciotti, Heidi Dierssen, Mark Dowell, John Dunne, Wayne Esaias, Bernard Gentili, Watson Gregg, Steve Groom, Nicolas Hoepffner, Joji Ishizaka, Takahiko Kameda, Corinne Le Quéré, Steven Lohrenz, John Marra, Frédéric Mélin, Keith Moore, André Morel, Tasha E. Reddy, John Ryan, Michele Scardi, Tim Smyth, Kevin Turpie, Gavin Tilstone, Kirk Waters, and Yasuhiro Yamanaka. A comparison of global estimates of marine primary production from ocean color. Deep Sea Research Part II: Topical Studies in Oceanography, 53(5-7):741–770, March 2006. ISSN 09670645. doi: 10.1016/j.dsr2.2006.01.028. URL https://linkinghub.elsevier.com/retrieve/pii/S0967064506000555.

[12] Marjorie A.M. Friedrichs, Mary-Elena Carr, Richard T. Barber, Michele Scardi, David Antoine, Robert A. Armstrong, Ichio Asanuma, Michael J. Behrenfeld, Erik T. Buitenhuis, Fei Chai, James R. Christian, Aurea M. Ciotti, Scott C. Doney, Mark Dowell, John Dunne, Bernard Gentili, Watson Gregg, Nicolas Hoepffner, Joji Ishizaka, Takahiko Kameda, Ivan Lima, John Marra, Frédéric Mélin, J. Keith Moore, André Morel, Robert T. O’Malley, Jay O’Reilly, Vincent S. Saba, Marjorie Schmeltz, Tim J. Smyth, Jerry Tjiputra, Kirk Waters, Toby K. Westberry, and Arne Winguth. Assessing the uncertainties of model estimates of primary productivity in the tropical Pacific Ocean. Journal of Marine Systems, 76(1-2):113–133, February 2009. ISSN 09247963. doi: 10.1016/j.jmarsys.2008.05.010. URL https://linkinghub.elsevier.com/retrieve/pii/S0924796308001176.

[13] V. S. Saba, M. A. M. Friedrichs, D. Antoine, R. A. Armstrong, I. Asanuma, M. J. Behrenfeld, A. M. Ciotti, M. Dowell, N. Hoepffner, K. J. W. Hyde, J. Ishizaka, T. Kameda, J. Marra, F. Mélin, A. Morel, J. O’Reilly, M. Scardi, W. O. Smith, T. J. Smyth, S. Tang, J. Uitz, K. Waters, and T. K. Westberry. An evaluation of ocean color model estimates of marine primary productivity in coastal and pelagic regions across the globe. Biogeosciences, 8(2):489–503, February 2011. ISSN 1726-4189. doi: 10.5194/bg-8-489-2011. URL https://bg.copernicus.org/articles/8/489/2011/.

